# Oral Feeding of Nanoplastics reduces Brain function of Mice by Inducing Intestinal IL-1β-producing Macrophages

**DOI:** 10.1101/2022.11.04.515261

**Authors:** Qianyu Yang, Huaxing Dai, Ying Cheng, Beilei Wang, Jialu Xu, Yue Zhang, Yitong Chen, Fang Xu, Qingle Ma, Ziying Fei, Fang Lin, Chao Wang

**Author notes:** Equally contribution (Qianyu Yang, Huaxing Dai, Ying Cheng). Corresponding authors: Chao Wang, 199 Ren’ai Rd. Suzhou, China, 86051265880722 Fan Lin, 199 Ren’ai Rd. Suzhou, China, 8613962509288.

## Abstract

**Background and Aims:** Nanoplastics (NPs) as contaminants in food and water have drawn an increasing public attention. However, little is known about how NPs shape the gut immune landscape after entering the body. The objective of the study was to explore indirect effects caused by the interaction of NPs with the mammalian gut and whole immune system after entering the body.

**Methods:** In this study, we fabricated NPs (∼500 nm) and microplastics (MPs) (∼2 μm) and aimed to evaluate their in vivo effects by feeding them in mice. The mechanism was then investigated by various technology including single-cell RNA sequencing of gut and brain tissue.

**Results:** The results suggested that NPs showed a better ability to induce gut macrophage activation than did MPs. In addition, NPs triggered gut interleukin 1 beta (IL-1β)-producing macrophage reprogramming via inducing lysosomal damage after phagocytosis. More importantly, IL-1β released from the intestine could affect brain immunity, leading to microglial activation and Th17 differentiation, all of which correlated with a decline in cognitive and short-term memory in NPs-fed mice

**Conclusions:** Thus, this study provides new insight into the mechanism of action of the gut-brain axis and delineates the way NPs reduce brain function, highlights the importance to fix the plastic pollution problem worldwide.

## Introduction

The global plastic pollution crisis is increasing, owing to the rapid growth in plastic production and use^1, 2^. Macroplastics that persist in the environment can break down into microplastics (MPs) (less than 5 mm) or nanoplastics (NPs) (less than 1 µm) after physical or chemical reactions. Micro or nanoplastics have been found in many food and drinking water, while there is no legislation for their concentration limitation in food. According to the World Wide Fund for Nature, an average person could ingest approximately 5 g of plastic every week^3^. In addition to MPs, NPs have attracted much attention owing to their size-dependent effect^4, 5^. The bioactivity of plastic particles increases with a decrease in particle size. This is because compared to large particles, small particles have a higher surface volume ratio and readily penetrate tissues and cells^6^. In addition to direct toxicity, the indirect effects caused by the interaction of NPs with the mammalian immune system after entering the body, associated with the consequences of perturbed immune responses in normal tissues, remain unclear.

Once plastic particles enter the body, the gut is the main site that immune cells recognize. The gut, as the body’s largest immune organ, plays an important role in regulation of the immune context^7^. Here we found that the induction of gut immune responses is inversely correlated with the size of orally-administered plastics. NPs are more capable than MPs in inducing and reprogramming macrophage activation. Using single-cell RNA sequencing technology, we observed a dramatic increase in the number of interleukin (IL)-1β^+^ macrophages in the intestine following NPs administration. More importantly, we unexpectedly found that compared to the control mice, long-term NPs-fed mice showed deficits in cognitive and short-term memory. These mental disorders are associated with the degradation of neurons and oligodendrocytes, activation of microglia, and differentiation of T cells to T helper 17 (Th17) cells in the brain, which are attributed to, at least in part, the IL-1β released from macrophages in the intestine. Thus, this study provides new insight into the mechanism of action of the gut-brain axis and delineates the way NPs reduce brain function, indicating the urgency of addressing plastic pollution within food globally.

## Results

### Characterization of plastic particles with different sizes

MPs and NPs were prepared by grinding relatively larger plastics. Most of the particles with irregularly shaped were observed under scanning electron microscopy (SEM) and transmission electron microscopy (TEM) (**Fig. 1a**). In line with the results of TEM, dynamic light scattering (DLS) analysis showed two kinds of plastic particles with an average diameter of ∼500 nm (NPs) and ∼2 μm (MPs) (**Fig. 1b**). In addition, four characteristic absorption peaks were observed in the infrared spectrum of the plastic particles (**Fig. 1c**), and the zeta potential of the plastic particles was approximately -10 mV (**Fig. 1d**), indicating that both MPs and NPs had similar chemical structures.

**Figure 1.**
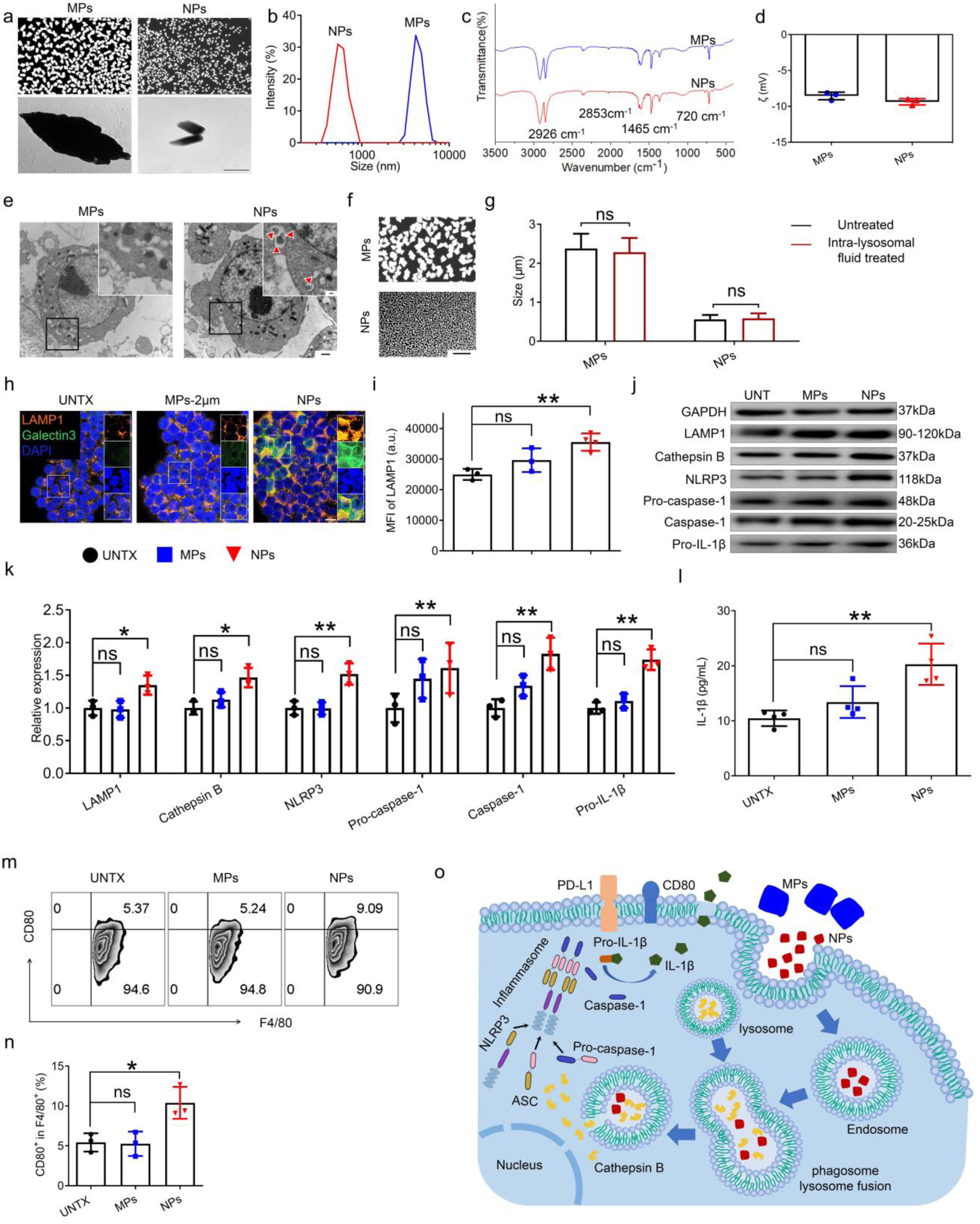
NPs are more likely than MPs to be ingested by macrophages and cause lysosomal damage. (**a**) SEM (above) and TEM image (below) of plastic particles of different sizes. Scale bar, 10 μm (above), 0.5 μm (below). (**b**) Particle size distribution measured by DLS. (**c**) Infrared spectrum of plastic particles with different sizes. (**d**) Zeta potential of three plastic particles in PBS (pH=7). (**e**) Bio-TEM image of plastic particles taken up by Raw264.7 cells. Scale bar, 0.2 μm (above) and 1 μm (below). (**f**) SEM images of plastic particles after treatment with lysosomal environment 12 hours and (**g**) Size distribution statistics of plastic particles before and after treatment. (**h**) Lysosomal damage revealed by confocal images of RAW264.7 cells. Red: lamp1, green: galectin3, blue: dapi. Scale bar, 10 μm. (**i**) Mean fluorescence intensity (MFI) of LAMP1 in Raw264.7 cells (gated on F4/80^+^). (**j**) Western blot analysis of the expression of various types of proteins in RAW264.7 cells after various treatments as indicated and (**k**) the relative expression of proteins compared to the untreated group (UNTX). (**l**) Cytokine IL-1β released from RAW264.7 cells after various treatments as indicated. (**m**) Flow cytometric analysis of CD80 expression on Raw264.7 cells after various treatments as indicated and (**n**) corresponding quantitative analysis. (**o**) Flow cytometric quantification of the MFI of PD-L1 in RAW264.7 cells after various treatments as indicated. (**p**) Schematic to explain the lysosomal damage pathway after cellular uptake of plastic particles. Data are shown as mean ± SD (n=3-4). Statistical significance was calculated by Student’s t-test (two-tailed) and one-way ANOVA using the Tukey posttest. \****P*** < 0.05; \*\****P*** < 0.01; \*\*\****P*** < 0.005; \*\*\*\****P*** < 0.001. a. u., arbitrary units.

Once plastic particles enter the body, they are primarily taken up by gut macrophages via endocytosis or phagocytosis. Endotoxin contamination was assessed in variety of particles used in the downstream cell and animal experiments (**Fig. S1a**). According to 3-(4,5-dimethylthiazol-2-yl)-2,5-diphenyltetrazolium bromide (MTT) and Annexin V/ PI double staining results, none of the MPs and NPs showed significant cytotoxicity to RAW264.7 cells at concentrations of 20 μg/mL within 24 hours (**Fig. S1b-d**). A high stability of plastic particles in cell culture medium was also confirmed (**Fig. S1e, f**). To investigate size-dependent phagocytosis of plastic particles, we incubated the MPs and NPs with RAW264.7 cells for 12 h. We used biological-TEM to observe cellular uptake. As shown in **Figure 1e**, NPs appeared to be taken up more effectively by macrophages than were MPs. In addition, the swallowed NPs were localized within lysosomal structures. We further examined whether the plastic particles could degenerate under lysosomal conditions. Plastic particles were incubated in an acidic solution (pH 5.5) for 12 h with hydrolytic enzymes, which simulated the internal environment of lysosomes. Qualitative and quantitative evaluations showed that the shape and size of the three types of plastic particles did not change significantly after treatment in a lysosomal environment (**Fig. 1f, g**), indicating their resistance to degradation within macrophages.

### NPs are more likely than MPs to be ingested by macrophages and cause lysosomal damages

As plastic particles are resistant to degradation in the lysosomal environment, we posit that lysosomal damage can be induced by the uptake of NPs. To determine this, we used galectin-3, which translocates to the membranes of leaky lysosomes, as a biomarker to study the potential lysosome damage induced by plastic particles^8^. Immunofluorescence images showed that the galectin-3 signal increased significantly in NPs-treated macrophages compared to MPs-treated macrophages (**Fig. 1h, Fig. S1g**). Lysosomal-associated membrane protein 1 (LAMP-1), a lysosomal marker, was also elevated in NPs-treated macrophages in comparison to MPs-treated ones (**Fig. 1i, j, Fig. S1h**). It is well established that defective lysosome degradation can cause lysosome enlargement^9^, leading to the enhancement of LAMP-1 levels. Damage to lysosomes leads to the release of substances that induce enhanced inflammatory signaling. The inflammasome/IL-1β axis is of particular interest, given its critical and unique role in macrophages ^10^. Western blot results showed that, compared to untreated or MPs treatment, NPs treatment significantly increased the levels of cathepsin B, nucleotide-binding oligomerization domain, leucine-rich repeat and pyrin domain-containing 3 (NLRP3), pro-cysteinyl aspartate specific proteinase (pro-caspase-1), caspase-1, and pro-IL-1β in the macrophages (**Fig. 1j, k, Fig. S1h**). As a result, there was a drastic increase in IL-1β levels produced by macrophages. In contrast, no difference was found in the MPs-treated macrophages, compared to the untreated control (**Fig. 1l**). The production of IL-6 and tumor necrosis factor-α (TNF-α) in the macrophages was similar after treatment in MPs and NPs (**Fig. S1i**). In addition, NPs increased the expression of CD80 in RAW264.7 cells, suggesting a cellular transition to a pro-inflammatory phenotype (**Fig. 1m, n**). To further corroborate that the cellular phagocytosis of NPs leads to IL-1β production, we used cytochalasin B which inhibits the phagocytic activity of macrophages, before incubation with NPs (**Fig. S2a-e**). These results demonstrated a direct correlation between NP phagocytosis and lysosome damage/IL-1β production by macrophages. More importantly, all the above data showed that, compared to MPs, NPs were more easily ingested into lysosomes by macrophages, which caused lysosomal damage, leading to increased production of IL-1β (**Fig. 1o**). We described a cascade of events in which NPs, rather than MPs, significantly induce IL-1β production by macrophages.

### NPs are more likely than MPs to perturb the intestinal ecosystem

To further explore how plastic particles influence the microenvironment *in vivo*, we orally administered BALB/c mice with the MPs and NPs (10 mg/kg) once daily for a week, the dosage of which was based on the daily dose of plastic particles ingested by humans^11, 12^ and previous animals studies^13^. Plastic particles were slightly degraded in the gastric fluid solution, indicating they could reach the intestine through the stomach (**Fig. S3a, b**). We observed that the length of the colon was shortened in MPs and NPs administrations compared with that in untreated mice. Moreover, the length of the colon was shorter in the NPs-treated group (**Fig. 2a, b**). Hematoxylin and Eosin (H&E) staining results showed that the intestine of NPs-treated mice had a marked disruption of the intestinal barrier and infiltration of immune cells, indicating the presence of colitis, which was not observed in MPs-fed mice (**Fig. 2c**).

**Figure 2.**
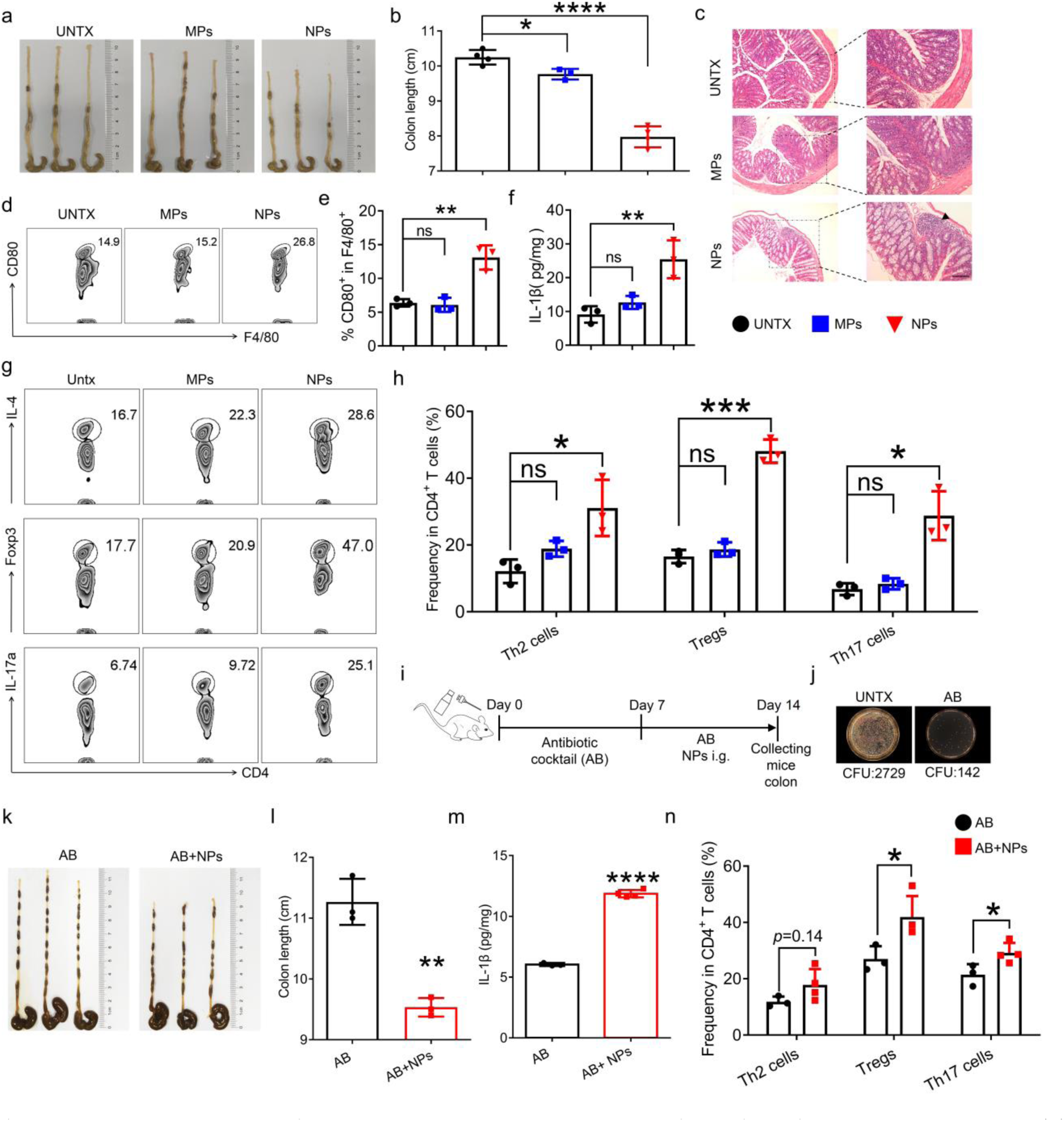
NPs are more likely than MPs to perturb the intestinal immune ecosystem. (**a**) Representative images of the colon of mice fed with three different sizes of plastic particles. (**b**) Quantification of the colon length. (**c**) Representative images of H&E-stained colon sections illustrate inflammatory cell infiltrates. Scale bars, 100 μm (left), 50 μm (right). (**d**) Flow cytometric analysis of CD80 expression on F4/80^+^ cells and (**e**) corresponding quantitative analysis in the intestine of mice fed with three different sizes of plastic particles. (**f**) The level of IL-1β in intestine measured by ELISA. (**g**) Representative plots and (**h**) percentages of IL-4^+^ (Th2 cells), Foxp3^+^ (Tregs) and IL-17a^+^ (Th17 cells) in CD4^+^ T cells in the mice fed with plastic particles of different sizes. (**i**) Scheme of mice treated with antibiotic cocktail (AB) before being fed with NPs. (**j**) Representative plot of bacterial colony growth after 24 hours of fecal coating of mice. (**k**) Representative images of the colon of AB-treated mice fed with NPs and (**l**) quantification of the colon length. (**m**) IL-1β in colon of AB-treated mice fed with NPs measured by ELISA. (**n**) Proportion of Th1 cells, Tregs and Th17 cells in CD4^+^ T cells in colon of AB-treated mice fed with NPs. AB: Antibiotic cocktail. Data are shown as mean ± SD (n=3-4). Statistical significance was calculated by Student’s t-test (two-tailed) and one-way ANOVA using the Tukey posttest. \****P*** < 0.05; \*\****P*** < 0.01; \*\*\****P*** < 0.005; \*\*\*\****P*** < 0.001. a. u., arbitrary units.

The immune context of the intestine was investigated. Consistent with the *in vitro* results, the number of macrophages associated with CD80 expression was significantly increased in the intestine of mice receiving NPs compared to that in the intestines of MP-treated or untreated mice (**Fig. 2d, e, Fig. S3c, d**). Macrophages are a major source of IL-1β ^14^. Among all three pro-inflammatory cytokines, the level of IL-1β was found to drastically increase in the intestine (**Fig. 2f**), as well as in the serum (**Fig. S4b**) following NPs treatment, compared to those in the control. Accumulating evidence suggests that macrophages play an important role in CD4^+^ T helper cell polarization. Further, using flow cytometry, we investigated the phenotype of CD4^+^ T cells in the intestinal ecosystem collected from mice after various treatments. In contrast to MPs-treated mice, those administered with NPs showed a marked increase in the populations of T helper 2 (Th2), Treg, and Th17 cells in CD4^+^ T cells (**Fig. 2g, h)**. We reasoned that Tregs and Th2 cells were induced in response to the inhibition of proinflammatory macrophages, whereas IL-1β secreted by macrophages directed and maintained Th17 polarization^15^.

Studies have shown that microplastics cause intestinal pathology by altering the gut microbiome^16^. We used antibiotic cocktail to indiscriminately eliminate intestinal microflora (**Fig. 2i, j**). In our experiment, antibiotic treatment did not lead to changes in the intestine obviously (**Fig. S6a-j**). Following this, the mice were administered the same dose of NPs for one week. Similarly, the length of the colon was shorter than that of the untreated mice (**Fig. 2k, l**). Intestinal IL-1β levels also significantly increased after NPs treatment **(Fig. 2m)**, as well as the alternation in immune context (**Fig. 2n, Fig. S7a-f**). Together, these results suggest that NPs are more likely than MPs to alter the intestinal ecosystem by directly interacting with macrophages, independent of the intestinal flora.

To further characterize and understand the intestinal immune microenvironment, we used 10× genomics of intestinal mononuclear cells isolated from mice receiving NPs for one week. Cellular markers identified five clusters of CD45^+^ cells, including B cells, T cells, granulocytes, plasmacytoid dendritic cells (pDCs), and macrophages (MΦ) &mast cells (**Fig. 3a, Fig. S8a-d**). We observed that mice administered NPs had an increased proportion of cells within MΦ &mast cells clusters as well as pDC clusters, while the frequency of T cells, B cells, and granulocytes did not change significantly (**Fig. 3b**). The violin diagram of gene expression of CD45^+^ cells showed that feeding of NPs up-regulated the expression of *Il1b* (IL-1β), *Lamp*1 (LAMP 1), and *Ctsb* (cathepsin B) (**Fig. 3c**), which indicated a significant enrichment in a secretory lysosomal pathway concentrated in the MΦ &mast cells subclusters (**Fig. 3d**).

**Figure 3.**
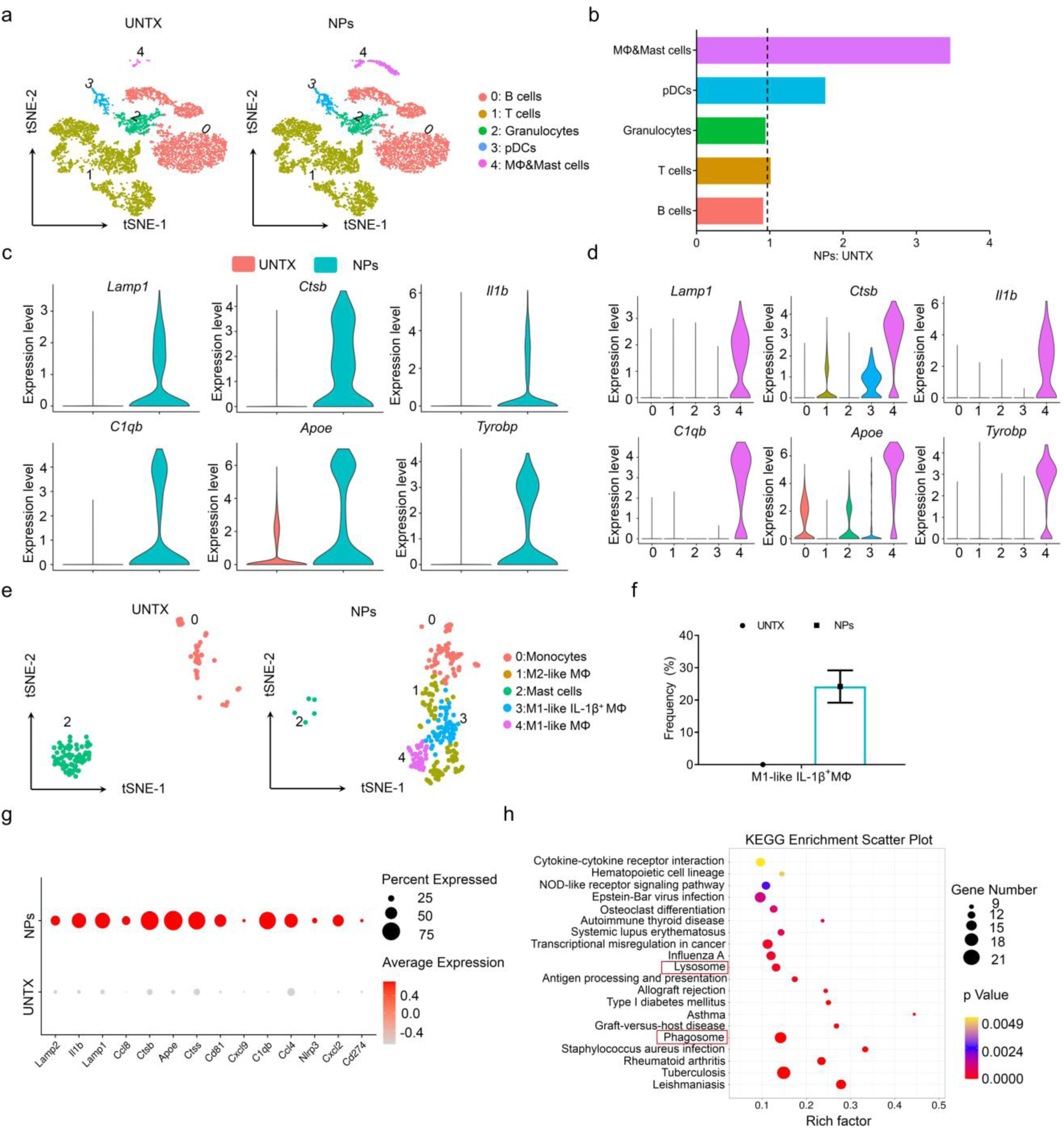
NPs alters the intestinal immune microenvironment. (**a**) tSNE plot of CD45^+^ cells from intestinal cells. Clusters identified as 5 different immune cells are indicated. (**b**) Relative proportions of each cluster within all clusters. (**c**) Violin plots of representative differential gene expression in all clusters and (**d**) the relative expression of the above genes in each cluster. (**e**) tSNE plot of MΦ & mast cells from CD45^+^ cells and (**f**) the proportion of IL-1β^high^ MΦ subcluster in this cluster. (**g**) Dotplot of differentially expressed gene in MΦ cells. (**h**) KEGG enrichment scatter plot of IL-1β^high^ MΦ subcluster.

We then focused on the MΦ cells subclusters to further divide them into subsets (**Fig. 3e**) with well-defined marker genes (**Fig. S9a, b**). The proportion of macrophages was significantly increased and that of mast cells decreased in mice administered NPs (**Fig. 3e**). We confirmed that *Il1b*-expressing immune clusters also expressed classical M1-like macrophage markers, including *Adgre1* and *Cd86*. Of note, we identified *Il1b*^+^ macrophages as a new cell cluster in NPs-treated mice (**Fig. 3f**), reinforcing that macrophages were major sources of IL-1β in the intestine of NPs-fed mice. In addition, we analyzed *Il1b* expression in all intestinal cells, including non-immune cells, and found its enrichment in myeloid cells (**Fig. S10**).

We then investigated the differentially expressed genes in macrophage subclusters. *Lamp1*, *Lamp2*, *Ctsb*, *Ctss* (cathepsin S), *Nlrp3,* and other genes related to the lysosomal damage pathway were significantly increased in NPs-fed mice compared with those in untreated mice (**Fig. 3g**). By analyzing the Kyoto Encyclopedia of Genes and Genomes (KEGG) pathway of *Il1b*^+^ macrophages, we also found that this cluster was closely involved in lysosome and phagosome-related pathways (**Fig. 3h**), indicating that IL-1β production was associated with lysosomal and phagosome damage. In summary, these data provide strong evidence that the oral administration of NPs induces high levels of IL-1β in the intestine by inducing lysosomal damage in macrophages.

### Long-term uptake of NPs drives alteration of brain immunity

In view of a recent study demonstrating that certain immune ecosystems in the gut can affect the brain, causing inflammation^17^, we evaluated whether feeding NPs could affect immunity in the brain. To test this, we studied the behavior of mice receiving NPs orally after two months. Institute of Cancer Research (ICR) mice and APP/PS1 mice were administered 10 mg/kg of NPs daily according to previous studies^11^. After two months, the behavior of the mice was tested, including open field behavior, the water maze test, and the novel object recognition test (**Fig. 4a**). To our surprise, the results of the open field experiment showed that the total locomotor activity of NPs-treated mice was less than that of the control mice and was more inclined to move out of the central region (**Fig. 4b-d, Fig. S11a-c**). The Morris water maze is a spatial learning test for rodents^18^. The swimming speed of the tested mice showed that motor function was not affected by feeding plastic particles (**Fig. 4e, f, Fig. S11d, e**). Unexpectedly, after one week of training, mice receiving NPs showed significantly more time to reach the target platform, indicating a reduction in the spatial memory of these mice (**Fig. 4e, g, Fig. S11d, f**). For new object recognition, there was no obvious difference between the two groups in the exploration of initial objects, but NPs-fed mice showed a significant decrease in the new object preference index in novel-object recognition, suggesting impairment of short-term cognitive memory capacity (**Fig. 4h, Fig. S11g**). Notably, the behavior of MPs-fed mice was similar to that of the control mice (**Fig. S12**). These results suggest that reduced mental functions in mice may be a consequence of long-term NPs uptake.

**Figure 4.**
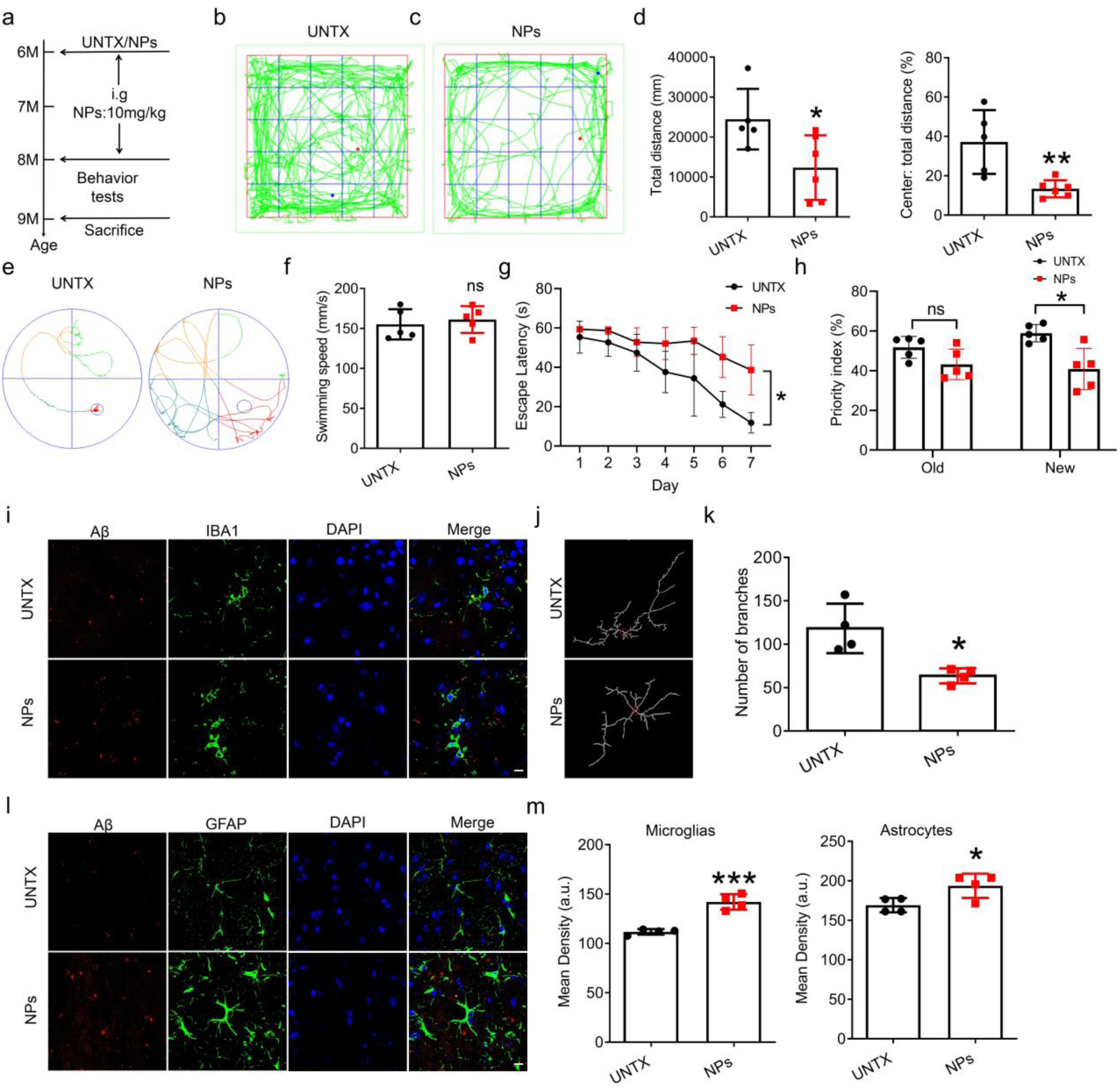
Long-term uptake of NPs drives abnormal of behavior in mice. (**a**) Experimental timeline. (**b**) Representative open field test images and (**c**) quantitative analysis of the total distance and (**d**) the ratio of the center distance to the total distance. (**e**) Plots of Morris water maze images and quantification of (**f**) the average swimming speed and (**g**) escape latency of day 1 to day 7. (**h**) Recognition index in the novel object recognition (NOR). (**i**) Confocal microscopy images of Aβ protein in microglia. (**j**) Plot of microglia branches and (**k**) corresponding quantitative analysis. (**l**) Confocal microscopy images of Aβ protein in astrocytes. (**m**) Mean fluorescence quantification of Aβ protein in microglia (left) and astrocytes (right). Red: Aβ, green: IBA1 or GFAP, blue: DAPI. Scale bar, 10μm (**i, l**). Data are shown as mean ± SD (n=4-6). Statistical significance was calculated by Student’s t-test (two-tailed) and one-way ANOVA using the Tukey posttest. *P < 0.05; **P < 0.01; ***P < 0.005; ****P < 0.001. a. u., arbitrary units.

We then want to study the underlying mechanism for this effect. It was found that H&E staining of the whole brain showed no obvious pathological features (**Fig. S13a**). Notably, we found that mice showed increased Aβ aggregation of the microglia and astrocytes in the hippocampus after long-term NPs treatment (**Fig. 5i-m, Fig. S13b, c**). Similar results were observed in the APP/PS1 mice (AD mice) (**Fig. S14**), indicating that long-term NPs treatment can exacerbate AD process. In addition, a decrease in the number of branches suggested synchronous microglial activation in NPs-fed mice (**Fig. 4j, k**). These results suggest that the long-term uptake of NPs drives alterations in the brain, which may contribute to the attenuation of cognitive memory capacity.

**Figure 5.**
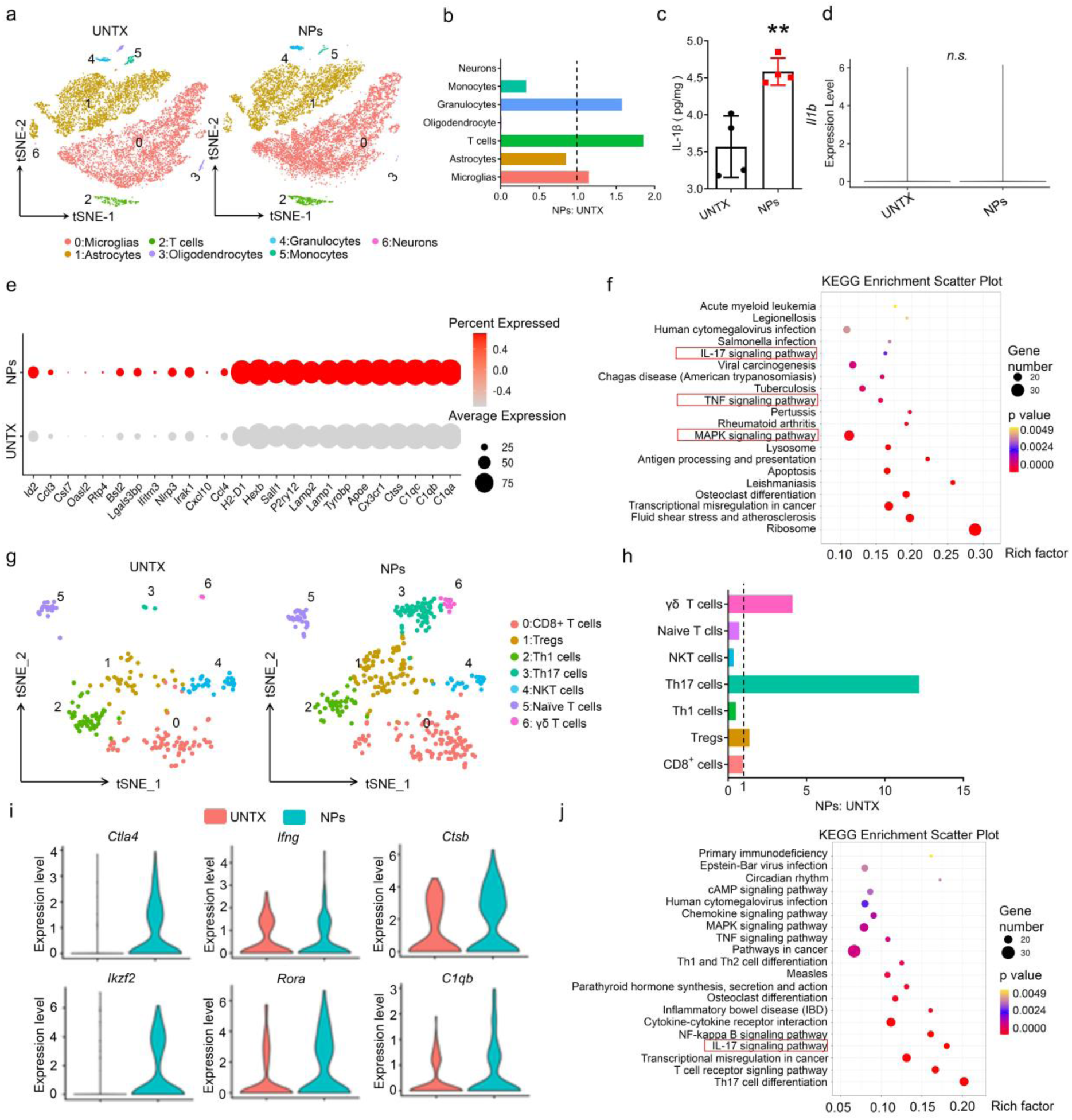
Long-term uptake of NPs drives alteration of brain immunity. (**a**) tSNE plot of all the brain cells and (**b**) the percentage of each cluster in total clusters. (**c**) Cytokine analysis of IL-1β in brain by ELISA. (**d**) violin diagram of *Il1b* expression in brain cells. (**e**) Dotplot of differentially expressed gene in microglia. (**f**) KEGG enrichment scatter plot of microglia. (**g**) tSNE plot of T cells from brain cells and (**h**) the percentage of each subset in this cluster. (**i**) Violin plots of representative differential gene expression in all subsets. (**j**) KEGG enrichment scatter plot of Th17 cells. Data are shown as mean ± SD (n=4-6). Statistical significance was calculated by Student’s t-test (two-tailed) using the Tukey posttest. \****P*** < 0.05; \*\****P*** < 0.01; \*\*\****P*** < 0.005; \*\*\*\****P*** < 0.001. a. u., arbitrary units.

We therefore reasoned that the underlying causes of Aβ formation likely due to the alteration of brain immunity. Further, we sought to determine the mechanism by which NPs influence brain immunity. We again performed a 10× single-cell map analysis of the whole brain of mice fed NPs for two months (**Fig. S15a**). We observed a significant increase in the number of microglia, granulocytes, and particularly T cells in the NPs-fed mice compared to those in the control mice (**Fig. 5a, b)**. In sharp contrast, a drastic decrease in the number and frequency of neurons and oligodendrocytes was observed in NPs-treated mice, suggesting that the functions of neurons in the brain were significantly affected. Previous studies have implicated IL-1β as an extracellular signal in the initiation of apoptosis in neurons and oligodendrocytes^19^. We then determined IL-1β in the brains of mice receiving NPs, which showed the IL-1β levels were significantly higher than that in untreated mice (**Fig. 5c**). Intriguingly, we did not find an IL-1β-producing microglia cluster of the brain using single-cell map analysis. In addition, the genetic violin diagram of *Il1b* revealed that *Il1b* was concentrated in brain granulocytes, while there was no difference in *Il1b* expression between untreated and NPs-treated mice (**Fig. 5d, Fig. S15b**). However, high IL-1β level had been observed in the NPs-treated intestine as well as in circulation, suggesting that the source of brain IL-1β is likely from the gut (**Fig. 5c, Fig. S16a, b**). Together, we reasoned that the degradation of neurons and oligodendrocytes was attributed to, at least in part, the IL-1β from the NPs-treated intestine^20^. Next, we identified many upregulated genes associated with immune activation, such as *Ccl3*, *Ccl4*, *Cx3cr1*, *Cxcl10* (chemokines); *Ifitm3*, *Oasl2*, *Rtp4*, *Bst2* (interferon-response genes); and *Lamp1*, *Lamp2*, *H2-D1*, *P2ry12*, *Hexb* (microglia activation-related genes)^22^. In addition, we found microglial clusters characterized by the increased expression of genes encoding the components of *Apoe,C1q (C1qa,C1qb,C1qc),* and *Tyrobp*, the upregulation of which has been reported in AD-related condition (**Fig. 5e**)^23^. KEGG pathway analysis of microglia showed that the cluster was involved in a variety of inflammatory pathways **(Fig. 5f**).

The ability of IL-1β to promote Th17 differentiation has been widely reported^24^. For T cells, we observed a drastic increase in the number of Th17 cells among CD4^+^ T cells, which was most likely induced by high levels of IL-1β (**Fig. 5g-h, Fig. S17**). Th17 has been implicated in the pathogenesis of many disorders with an inflammatory basis, including neurological diseases^15^. Expression of the Th17 proinflammatory cytokine IL-17a, exacerbates Aβ deposition and increases neurodegeneration^25^. In NPs-fed mice, increased IL-17a level were found in the brain (**Fig. S18**). More importantly, we identified *Apoe*, *C1qb*, and *Tyrobp*, whose expressions were related to AD^26^, were all upregulated in the brain of NPs-fed mice (**Fig. 6i**). Next, KEGG enrichment analysis of Th17 clusters showed significant Th17 cell-mediated neuroinflammation were induced in the brains of NPs-treated mice (**Fig. 5j**). This evidence suggests a link between NPs feeding and brain immunity alteration mediated by IL-1β signaling.

**Fig. 6.**
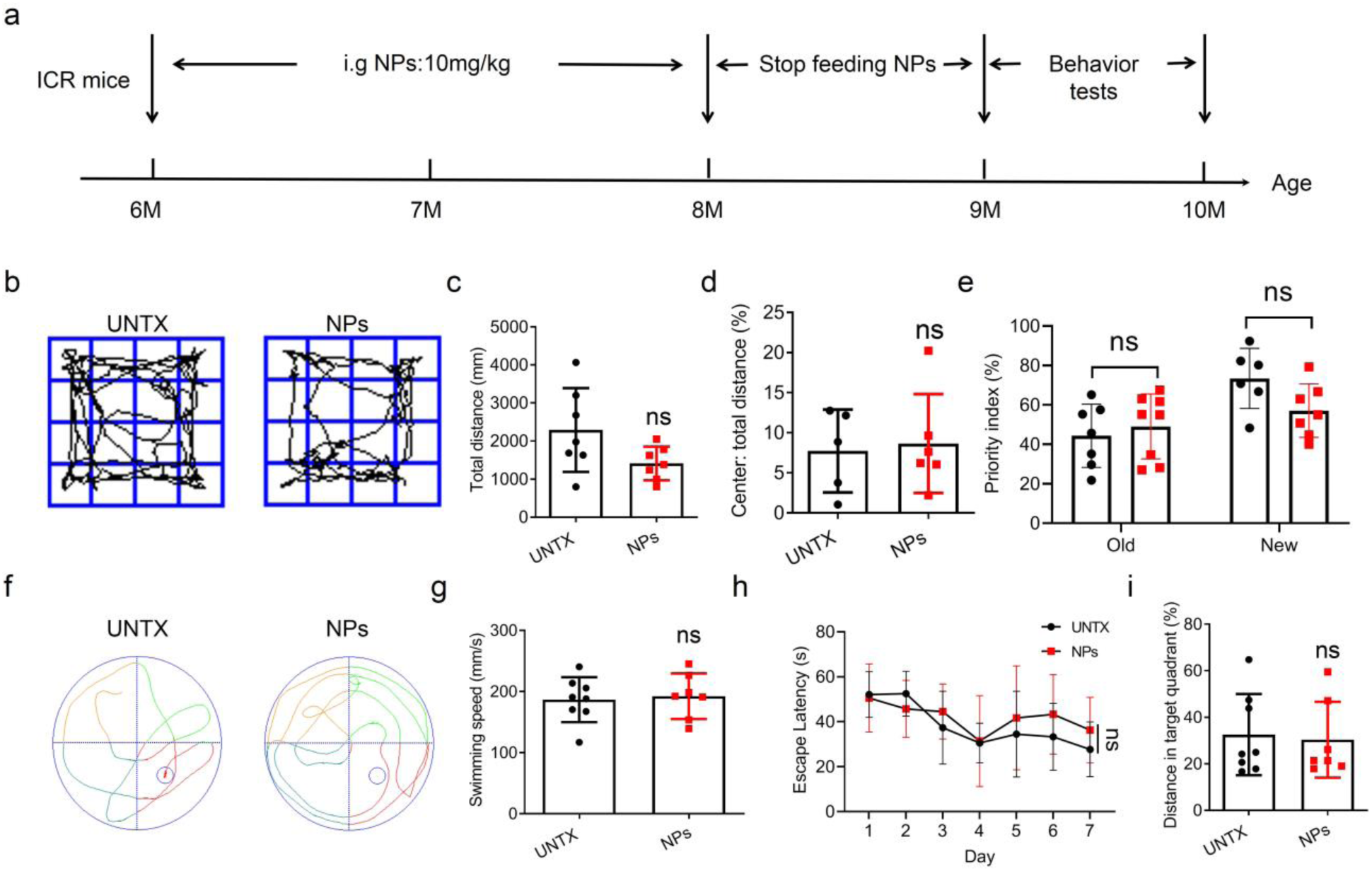
Discontinuation of NPs feeding alleviates behavioral disorders caused by NPs. (**a**) Experimental timeline. (**b**) Representative open field test images and (**c**) quantitative analysis of the total distance and (**d**) the ratio of the center distance to the total distance. (**e**) Recognition index in the novel object recognition (NOR). (**f**) Plots of Morris water maze images and quantification of (**g**) escape latency, (**h**) swimming speed, and (**i**) the ratio of the distance in the target quadrant to the total distance. Data are shown as mean ± SD (n=6-8). Statistical significance was calculated by one-way ANOVA using the Tukey posttest. \****P*** < 0.05; \*\****P*** < 0.01; \*\*\****P*** < 0.005; \*\*\*\****P*** < 0.001.

### Anti-IL-1β mAb effectively alleviates behavioral barrier caused by NPs

To investigate whether neutralizing IL-1β would rescue the brain damage caused by NPs feeding, we intraperitoneally injected anti-IL-1β mAb (α-IL-1β, 10 mg/kg, twice a week) into NPs-treated mice. After two months, the behavior of the mice was tested. The results showed that α-IL-1β treatment effectively alleviated abnormal behavior of NP-fed mice (**Fig. S19**). However, the effects of α-IL-1β treatment were not observed in AD (APP/PS1) mice (**Fig. S20**), which warrants further investigation. These results confirmed that IL-1β paled a key role in inducing cognitive and memory impairment.

### Discontinuation of NPs feeding alleviates behavioral disorders caused by NPs

Finally, we questioned whether stop NPs feeding alleviate the effects of brain damage. After two months NPs treatment, we stopped NPs feeding for one month, then the behavior of the mice was tested (**Fig. 6a**). Encouragingly, the results showed that the reduced motor activity and the new object exploration priority index induced by NPs treatment in mice was alleviated after discontinuation of NPs feeding (**Fig. 6b-e**). For the Morris water maze, there was no difference between the two groups in the time to reach the target platform, and the probability in the target quadrant (**Fig. 6f-i**). In addition, the effects of this strategy of discontinuing NPs feeding also observed in AD (APP/PS1) mice (**Fig. S21**). These evidences strongly suggested that the cognitive dysfunction caused by long-term NPs treatment is reversible rather than permanent, if the mice were stop exposure to NPs for at least one month.

### Melanin rescues the inflamed intestinal environment

Previous reports have shown that melatonin can effectively alleviate the inflammatory response induced by IL-1β^27, 28^. We hypothesized that treatment with melatonin may rescue the inflamed intestinal microenvironment established by the oral administration of NPs. To evaluate this, mice were fed NPs (10 mg/kg daily for one week) and administered a dose (10 mg/kg) of melatonin orally on days 1, 3, and 5. The results showed that melatonin treatments significantly alleviated the shortening of colon length (**Fig. 7a, b**) and pathological lesions (**Fig. 7c**) caused by NPs. Flow cytometry analysis showed that the ratio of CD80^+^ F480^+^ macrophages in the intestine of melatonin-treated mice decreased (**Fig. 7d**), with reduced pro-inflammatory cytokine, particularly IL-1β, generation both in the intestine and in serum (**Fig. 7f, g**). In addition, the proportion of Th17 cells in CD4^+^ T cells and Treg cells decreased significantly (**Fig. 7h, i**), accompanied by a decrease in IL-17a content in the intestine and serum. In summary, these results showed that melatonin treatment reversed the microenvironment in the intestine of NP-fed mice.

**Figure 7.**
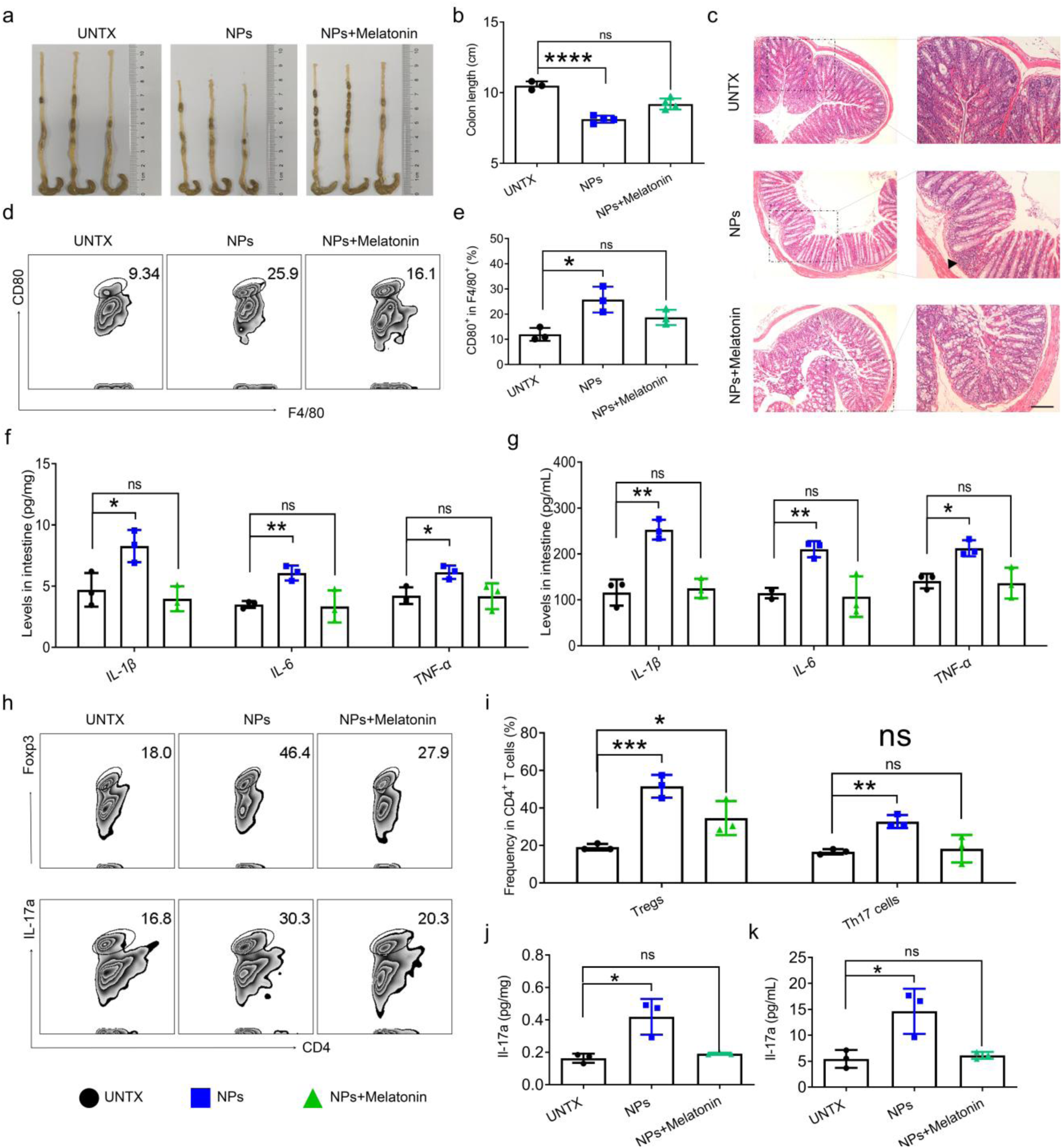
Melatonin rescues the inflamed intestinal environment. (**a**) Representative images of the colon of mice after various treatments as indicated and (**b**) quantification of the colon length. (**c**) Representative H&E images of the intestine of mice after various treatments as indicated. Scale bars, 100μm (left), 50 μm (right). (**d**) Representative plots of flow cytometric analysis of CD80 expression on F4/80^+^ cells in the intestine and (**e**) corresponding quantitative analysis. (**f**) Cytokine analysis of IL-6, TNF-α, and IL-1β in intestine and (**g**) in serum of mice after various treatments as indicated. (**h**) Representative plots showing the fraction of Foxp3^+^ and IL17a^+^ cells in CD4^+^ T cell population in the intestine after different treatments as indicated and (**i**) corresponding quantification results. (**j-k**) Cytokine analysis of IL-17a (**j**) in intestine and (**k**) in serum of mice after various treatments as indicated. Data are shown as mean ± SD (n=3-4). Statistical significance was calculated by one-way ANOVA using the Tukey posttest. \****P*** < 0.05; \*\****P*** < 0.01; \*\*\****P*** < 0.005; \*\*\*\****P*** < 0.001.

## Discussion

In our study, we found that although the NPs were effectively engulfed by macrophages, they were resistant to degradation within their lysosomes. As a result, a cascade of events is triggered by lysosomal damage, culminating in the reprogramming of IL-1β-producing macrophages. In NPs-fed mice, our unbiased single-cell sequencing identified newly clustered IL-1β^+^ macrophages, which release a large amount of IL-1β form gut to the whole body.

We identified IL-1β as a mediator in the cross-talk between the gut and brain, causing neuronal death, microglial activation, and Th17 differentiation. After long-term feeding with NPs, we observed increased levels of IL-1β in the both intestine and brain of the mice. More importantly, we did not find an IL-1β-producing microglia cluster in the hippocampus of the brain using single-cell map analysis, suggesting that the source of brain IL-1β is more likely from the gut. In addition, Dunn et al. indicated that peripheral administration of IL-1α or IL-1β in low doses can alter behavior.^29^ Our study provides new insights into the mechanism of action by which the gut communicates with the brain through cytokine signaling.

Studies have shown that MPs cause intestinal pathology by altering the gut microbiome^16, 30^. However, our study, did not recognize any overriding influence of the gut microbiome in manipulating the intestinal immune microenvironment in NPs-fed mice. IL-1β-producing macrophages appeared even after eliminating most of the gut microbes using antibiotics, suggesting that NPs are more likely to alter the intestinal ecosystem by interacting with macrophages directly. However, we cannot rule out the possibility that the gut microbiome regulates gut immunity in NPs-fed mice. Many studies have indicated that gut vascular barrier leakiness and dysfunction during enteritis lead to bacterial translocation into circulation and other organs, including the brain.^21^ The influence of NPs on the gut microbiome and potential bacterial translocation from the intestine to other organs needs to be investigated in future studies.

A recent study revealed that the death of neurons is before Aβ formation and neurodegenerative disease^31^. We therefore reasoned that the underlying causes of Aβ formation likely due to the activated microglia that impair generation and induce the death of myelin-forming oligodendrocytes and neurons in the hippocampus, and IL-1β cytokine from gut contributes to at least part of this dysregulation. This raises a worrying speculation that long-term exposure to NPs may increase the risk of developing AD. We also discovered that the cognitive dysfunction caused by long-term NPs treatment is reversible rather than permanent. Our study showed one more reason why we need to fix the plastic pollution problem.

These data suggest that IL-1β may be a therapeutic target. Behavioral impairments in mice were alleviated by an intraperitoneal injection of anti-IL-1β mAb. In addition, we have also proposed interventions to mitigate intestinal damage caused by NPs using immunomodulatory drugs (melatonin). We have demonstrated that the administration of melatonin rescued the inflamed microenvironment *via* reprogramming the intestinal immune milieu, thereby providing an effective low-cost approach to reduce potential damage in the brain during long-term exposure to NPs. Other clinically approved anti-inflammatory agents can be examined in the future.

In conclusion, our study identified that gut IL-1β-producing macrophages could be induced by the oral administration of NPs rather than MPs. IL-1β-producing macrophages in gut affect brain immunity by inducing microglial activation and Th17 differentiation, leading to mental deficits in NPs-fed mice. Our work provides new insights into the mechanism of the gut-brain axis and delineates that NPs induce a reduction in cognitive and memory abilities, highlight the importance to fix the plastic pollution in the food and drinking water problem worldwide.

## Methods

### Animals

BALB/c mice (6-month-old) were obtained from the Experimental Animal Center of Soochow University (Suzhou, China). APP-PS1 mice (6-month-old, half male and female) and ICR mice (6-month-old, half male and female) were purchased from Changzhou Cavens Experimental Animal Co. Ltd (Suzhou, China). The mice were housed in stainless-steel cages and acclimated for two weeks at 21–24 °C, 50–55 % relative humidity, and a 12 h light/dark cycle. Mice were not fasted. Food and water were provided *ad libitum*. All animals were tested under ethically compliant supervision of the ethics committee of Soochow University (approval number: SUDA20210922A01).

We used the ARRIVE reporting guidelines. The objective of the study was to explore indirect effects caused by the interaction of NPs with the mammalian immune system after entering the body, associated with the consequences of perturbed immune responses in brain tissues. we orally administered BALB/c mice with the MPs and NPs (10 mg/kg) once daily for a week to two months, the dosage of which was based on the daily dose of plastic particles ingested by humans^11, 12^ and previous animals studies^13^. According to the World Wide Fund for Nature, an average person could ingest approximately 5 g of plastic every week (through formula conversion [human equivalent dose (mg/kg) = mouse dose (mg/kg) × (mouse Km/Human Km)^0.33^)^3^. The experimental group sizes were approved by the regulatory authorities for animal welfare, after being defined to balance statistical power, feasibility, and ethical aspects. Animals were randomly assigned to groups based on body weight, using a computer based random order generator. All healthy animals were included in the study. No animals were excluded in this study. The investigators were not blinded to allocation during experiments and outcome assessment. Animals were euthanized when exhibiting signs of impaired health. All experiments were run at least in triplicate.

## Method

### Cell culture

The mouse mononuclear macrophage leukemia cell line RAW264.7, and the BALB/c mouse-derived colorectal carcinoma cell line CT26-Luc were obtained from the American Type Culture Collection. RAW264.7 cells were cultured in low-glucose Dulbecco’s Modified Eagle Medium (Corning, Wujiang, China) containing 10 % fetal bovine serum (FBS) (Bovogen, Melbourne, Australia), penicillin (100 U/mL; Invitrogen, New York, USA), and streptomycin (100 U/mL; Invitrogen, New York, USA). For routine maintenance in culture, RAW264.7 cells were seeded at a confluence of approximately 5–10 % (1×10^5^ cells in a 100 mm petri dish) and grown to a confluence of approximately 20–30 % (3×10^6^ cells in a 100 mm petri dish). For cell passaging, the cells were easily separated from the dish wall by repeated blowing using pipettor (Eppendorf, Hamburg, Germany). CT26-Luc cells were maintained in Roswell Park Memorial Institute 1640 medium (Corning, Wujiang, China) containing 10 % FBS (Bovogen, Melbourne, Australia), penicillin (100 U/mL; Invitrogen, New York, USA), and streptomycin (100 U/mL; Invitrogen, New York, USA). During cell passaging, the CT26-Luc cells were digested using trypsin (HyClone, Utah, USA) for 3 min before collection. All solutions, buffers, and media were prepared using sterile, distilled, and deionized water. The temperature of the incubator was maintained at 37 °C, and the CO_2_ concentration was 5 %.

### Preparation of different particles

PE plastic was purchased from Guangdong Huachuang Plasticizer (Guangdong, China), and MPs were prepared by grinding relatively larger plastics. For *in vivo* experiments, NPs and MPs were suspended in a 0.5 % carboxymethylcellulose solution (CMC), wherein CMC was used as a co-solvent and sonicated for 10 min prior to administration. For *in vitro* experiments, the plastic particles were dissolved in phosphate buffered saline (PBS). The NPs samples were tested to confirm the absence of endotoxin contamination. As the polyethylene particles were light in cell culture media, submerged exposure of NPs to macrophages was realized by reducing the volume of cell culture media in cell culture dishes (40 μL/well of 96-well cell culture plates, 150 μL/well for 24-well cell culture plates). RAW264.7 cells (1×10^6^ cells) contained 20μg/mL plastic particles in cell culture media were added in petri dish Cetyltrimethylammonium bromide (CTAB) 2g and TEA 320 mg were weighed, 100 mL pure water was added, stirred at 96℃ and 500 RPM, and ethyl teosilicate was added with microfluidic device, and the reaction lasted for 45 min. After the reaction, supernatant was centrifuged. Wash with pure water at 15500 RPM for 3 times, 45 minutes, and then with ethanol for 3 times. Subsequently, ethanol and concentrated hydrochloric acid were added and reflux was performed for 24 h to remove excess CTAB. Centrifuge at 15500 RPM for 45 minutes, discard the supernatant, and wash the precipitate with pure water for three times. Finally, SiO2 NPs was obtained by drying at 60-80℃. TiO2(Macklin, catalogue no. T818936, Shanghai, China) purchased from Shanghai Maclean Biochemical Co., LTD. The preparation of SiO_2_ and TiO_2_ was described above.

### Characterization of dfifferent particles

The three types of plastic particles (MPs and NPs) were dispersed in double-distilled water and characterized by SEM and transmission electron microscopy TEM. In detail, the carbon-coated 400-mesh grid was initially activated and the plastic particles were then dropped onto the grids. Finally, the plastic particles were dried and observed using TEM. For SEM, we used double sticky carbon tape to deposit a silicon wafer dripped with plastic particles on the sample stage and then dried it overnight. DLS was used to measure the size of the plastic particles. The surface potential of the plastic particles was analyzed using a zeta potential analyzer. To determine the chemical properties of the plastic particles, an fourier-transform infrared spectrometer (V70 & Hyperion1000) was used to detect their characteristic peaks.

### Calculation of the number of particles

Based on the SEM and TEM results, we considered that the volume of plastic particles is approximately proportional to the cubic power of the particle size (r). The calculation of the number of particles is based on the ellipsoid volume calculation formula: volume (V)= 4πr^3^/3 and mass (m)= density (ρ, 0.96g/cm^3^)×V, which gives the mass per particle approximately. The number of particles was obtained from the total mass of the MPs and NPs used in the experiments.

### Cellular uptake of plastic particles

Cellular uptake of plastic particles was observed using a biological TEM. As the PE particles were light in weight in the cell culture media, submerged exposure of macrophages to NPs was achieved by reducing the volume of cell culture media in cell culture dishes (40 μL/well of 96-well cell culture plates, 150 μL/well for 24-well cell culture plates). Briefly, RAW264.7 cells (1×10^6^ cells) containing 20 μg/mL plastic particles in cell culture media were added to a petri dish. After co-incubation for 12 h, the supernatant was removed, the residual material was washed with PBS, the cells were collected in a centrifuge tube, and 2.5 % glutaraldehyde solution was added for fixation. After 24 h, samples were embedded and sectioned. The fixative was then removed and the samples were rinsed with PBS thrice for 15 min each time. Next, the samples were fixed with 1 % osmium solution for 1–2 h. The osmium waste solution was carefully removed and the samples were rinsed thrice with PBS for 15 min each. The samples were dehydrated with ethanol solutions of gradient concentrations (including six concentrations of 30, 50, 70, 80, 90, and 95 %) for 15 min each, and then treated with 100 % ethanol for 20 min; finally, over to pure acetone for 20 min. The samples were treated with a mixture of encapsulant and acetone 1:1 v/v for 1 h, 3:1 v/v for 3 h, and finally treated with pure encapsulant overnight. The permeation-treated samples were embedded and heated at 70 °C overnight to obtain the embedded samples. The samples were sliced in an ultrathin slicer (LEICA EM UC7, Leica, Wetzlar, Germany) to obtain 70–90 nm sections, which were stained with lead citrate solution and 50 % ethanol-saturated uranyl acetate solution for 5–10 min each, dried, and observed using TEM.

### *In vitro* cell cytotoxicity assay

The cytotoxicity of the conjugates was determined using MTT cell proliferation assay, according to the manufacturer’s protocol. Briefly, RAW264.7 cells (1×10^4^ cells/well) were plated in a 96-well plate. After the cells adhered to the wall, the supernatant was removed, and 40 μL of medium containing the three types of plastic particles was added at the set concentrations. After an additional 24 h of incubation, 5 mg/mL MTT in PBS (20 μL) was added to each well and incubated for another 4 h. Finally, the medium was replaced with 100 μL dimethylsulfoxide to dissolve the purple formazan crystals. After sufficient dissolution, the absorbance of the solution was measured at 570 nm using a multifunction microplate reader.

### Lysosomal damage

Small discs were placed in 24-well plates in advance; RAW264.7 cells were incubated with the three kinds of plastic particles in the above method for 24 h, and then the supernatant was removed. After washing with PBS, 4 % paraformaldehyde was added to fix the cells. After 30 min, the supernatant was removed and FACS buffer (PBS containing 3 % bovine serum albumin) was added after PBS cleaning and blocked for 15 min. PE-lamp-1 and FITC-galectin-3 were then added and incubated overnight in the dark. The following day, the cells were washed thrice with PBST (PBS containing 0.05 % tween20), stained with 4′,6-diamidino-2-phenylindole dye for 10 min to label the nuclei, and photographed using a confocal microscope (Zeiss LSM 800, Oberkochen, Baden-Württemberg, Germany).

### In *vitro* and *in vivo* simulation environment

A pH 5.5 acidic solution was prepared to simulate the intracellular lysosomal environment, and various hydrolytic enzymes, including acid phosphatase, nuclease, cathepsin, glycosidase, lipase, phosphatase, and sulfatase were added. In addition, artificial gastric fluid (Leagene, Beijing, China) containing pepsin and hydrochloric acid was used to simulate the gastric environment of the plastic particles after they entered the body. Briefly, plastic particles were incubated in both solutions for 12 h and their morphology was observed using an SEM.

### Western Blotting and cytokine detection

The RAW264.7 cells treated with the scheme mentioned above were collected, and a mixture of radioimmunoprecipitation assay lysate and protease inhibitor (50:1) was added. After cryo-lysis for a period of time (30-40 min), the extracted protein was quantified using bicinchoninic acid protein quantitative kit (Beyotime, Shanghai, China). The protein was denatured at high temperature, after which 1 × loading buffer was added, and the protein was separated using 10 % sodium dodecyl-sulfate polyacrylamide gel electrophoresis. The protein was then transferred to a polyvinylidene difluoride membrane by wet transfer, blocked using 3 % FACS buffer, and co-incubated with glyceraldehyde 3-phosphate dehydrogenase, anti-LAMP1, anti-cathepsin B, anti-NLRP3, anti-Pro-caspase-1, anti-caspase-1, anti-Pro-IL-1β, and anti-IL-1β (1:1000, applicable to each of these antibodies). Horseradish peroxidase-conjugated goat anti-rabbit immunoglobulin G secondary antibody (1:5000) was then incubated with the primary antibody. The hybridization bands of the target protein were detected using fluorescence chemistry, and quantitative analysis was performed using ImageJ software.

The cell supernatant of RAW264.7 cells was collected after 24 h of co-incubation with plastic particles. Mouse tissues in the *in vivo* experiment were lysed to obtain the tissue homogenate, and the supernatant was obtained using centrifugation. The cytokine content in the solution was determined using an enzyme-linked immunosorbent assay kit, according to the manufacturer’s instructions.

### Antibiotic cocktail treatment

Mice were provided autoclaved drinking water supplemented with a cocktail of the following broad-spectrum antibiotics as previously described: ampicillin (1 mg/mL, Solarbio, Beijing, China), gentamicin (0.5 mg/mL, Aladdin, Shanghai, China), metronidazole (1 mg/mL, Energy Chemical, Anqing, China), and vancomycin (0.25 mg/mL, Biosharp, Shanghai, China)^39, 40^. Antibiotic treatment was started one week before the experiments and continued for the duration of the experiments. To assesses the effect of the treatment, we used fresh feces from mice and performed dilution coating of the plates. The diluent was sterilized with double distilled water. All of the above processes were carried out on an ultra-clean platform, and the equipment used in the experiment was sterilized. After incubation in agarose medium for 48 h, fecal samples were counted using a colony counter (Czone 8, Hangzhou, China). Colony-forming units were used as the quantitative units for plate counts.

Immune indices of plastic particles were analyzed by continuous administration to mice at a dose of 10 mg/kg/day. The plastic particles were prepared as described above and suspended in 0.5 % CMC to obtain a 1 mg/mL suspension. In the experiment, the suspension was administered by gavage, and the volume of the gavage for each mouse was approximately 200 μL. After a week or two months of treatment, the organs of the mice were isolated and tissue homogenates were obtained using a tissue fragmentation machine. The tissue homogenates were then filtered using a nylon gauze to obtain a single-cell suspension. For flow cytometric immunoassay, single-cell suspensions were washed twice with PBS, centrifuged, and suspended in FACS buffer (PBS containing 1 % FBS). Further, the cells were stained with a fluorescent antibody (BioLegend, California, USA). The stained cells were analyzed using a BD Accuri C6 flow cytometer (BD Biosciences, Franklin Lakes, New Jersey, USA). For cytokine detection, 200 μL cell lysate was added to the single-cell suspension, incubated at 4 °C for 20–30 min, centrifuged at 12000 rpm for 10 min, and the supernatant was collected for subsequent detection. In addition, before removal of the organs from the mice, fresh blood was obtained by eyeball extraction and placed in a microfuge tube. After standing at 4 °C for 1 h, serum was obtained by centrifugation at 12000 rpm for 15 min, and was subsequently analyzed.

### Histological analysis and immunohistochemical staining

At the end of the experiments, the organs were fixed in 4 % neutral-buffered paraformaldehyde and embedded in paraffin. Paraffin sections were cut into 4 mm thick slices for H&E staining. For immunohistochemical studies, tissues were harvested, fixed in 4 % neutral buffered paraformaldehyde, embedded in paraffin, and prepared into serial sections (6 μm). Subsequently, the sections were dewaxed and rehydrated with xylene, ethanol, and deionized water. Endogenous peroxidase was quenched with 3 % hydrogen peroxide for 0.5 h and rinsed with deionized water, followed by blocking with 10 % normal goat serum for 1 h. Next, the slides were stained with rabbit anti-goat overnight at 4 °C, followed by incubation with horseradish peroxidase-conjugated secondary antibody for 1 h at room temperature (25 °C). Diaminobenzidine was used as the substrate to produce an observable brown color. Of note, considering that behavioral experience itself modifies brain, the brain histological analyses and immunohistochemical staining were performed in behaviorally naïve animals.

### Behavioral tests

The behavioral tests involved the following three experiments: the open field test, new object recognition test, and Morris water maze test. The open field test was performed first, followed by the new object recognition test on the second day, and finally the water maze experiment. Experiments were conducted on different days using the same mouse. Untreated mice were tested first, followed by experimental mice. The mice in the same group were tested according to their serial numbers. None of the behavioral tests changed from the beginning to the end of our study. The following parameters were assessed: Priority recognition index, quantitative analysis of the total distance and the ratio of the center distance to the total distance, quantification of the average swimming speed and escape latency. Notably, considering that behavioral experience itself modifies the brain, brain histological analyses were performed in behaviorally naïve animals.

#### Open field test

The assessment was conducted in a square white box. The mice were placed in the arena and allowed to move freely for approximately 10 min while being recorded by an overhead camera. Between tests, the box was cleaned with 75 % ethanol and dried. The footage was then analyzed using an automated tracking system for the following parameters: distance moved, velocity, and time spent in the predefined zones. Activity levels were assessed by calculating the total distance moved by the mice and the distance in the central area/total distance.

#### Novel object recognition test

On the first day, the mice were placed in an open-field (OF) box consisting of a quadrangular area (60 cm×60 cm×60 cm) and were habituated for 15 min. The next day, the training trial was performed by presenting the mice with two similar objects (A1 and A2, two identical yellow dice) for a period of 5 min. The testing trial was performed for 4 h; one of the two objects (A2) was replaced with a new object (B, red triangle), and the animals were left in the OF box for 5 min. Exploration of the objects was considered when sniffing or deliberate contact occurred with the objects or when the animal’s snout was directed toward the object at a distance of <1 cm. In addition, the cages were cleaned and dried using 75 % ethanol. The amount of time required to explore a new object provides an index of the recognition memory. The exploration time for familiar or novel objects during the test phase was recorded by an observer who was blinded to the group of mice. Priority recognition index = time to explore A2 or B/ (time to explore A1+ time to explore A2 or B) ×100 %.

#### Morris water maze test

The pool was equally divided into four invisible quadrants, and a hidden platform was set in one quadrant submerged 1 cm beneath the water surface. Several colorful signs (a triangle, a circle, a square, and a five-pointed star designed by a construction paper) were placed in each quadrant around the pool, which acted as clues for mice during the test. The inner wall of the pool was pasted with white wallpaper and a small amount of TiO_2_ (food grade) was used to make the water opaque. The pool temperature was maintained at 22 °C using a temperature controller. A camera with a digital tracking system was fixed over the pool to automatically identify the swimming track of the mice (swimming track from the starting point to the platform). The experimental period was 7 days, including a hidden platform test (days 1–6) and probe test (day 7). During the hidden platform test, the mice were randomly released into the water from each quadrant to explore and find the hidden platform by themselves for 60 s. The escape latency, defined as the time for the animal to find the platform, was recorded from day 1 to day 7. Mice were tested in two different quadrants per day at an interval of 4 h, and their latency was recorded. To examine spatial reference memory, a probe test was performed 24 h after the last day of the hidden-platform test. During the probe test, the platform was removed from the pool and the mice were allowed to swim freely for 60 s. Similarly,, the tracks were recorded using an overhead camera linked to a computer for visualization purposes. The swimming speed of the tested mice was determined to preclude motor function deficits as a confounder. To ensure that the olfactory cues of the previous group of mice in the water maze were removed, the water in the maze was changed between tests.

### Intestinal permeability assay to dextran molecule

NP-treated and untreated mice were perfused with FITC-dextran (MW 4000, MCE) by gavage, and whole blood was collected after 4 h. The blood samples were placed in a 4 °C refrigerator for 1 h and centrifuged at 5000 rpm for 15 min to collect serum. Serum (100 μL/well) was added to a black fluorescent plate, and the absorbance value at 490/525 nm was measured using a microplate reader (BioTek, Beijing, China). Higher serum dextran levels represent greater intestinal permeability.

### Antibody neutralization

During NPs treatment, mice were injected intraperitoneally with 10 mg/kg anti-mouse/rat IL-Iß (BioXCell, New Hampshire, USA) twice a week. Two months later, the mice underwent behavioral tests, including the open field, novel object recognition, and Morris water maze tests.

### Single cell RNA sequencing of the intestine and brain

Single-cell RNA sequences of the colon and brain were obtained from untreated or NPs-treated mice for 7 days and 2 months, respectively. The tissues were enzymatically digested and processed into a single-cell suspension. The oligo-dT-based complementary DNA database is a droplet-partioning barcode using the Chromium Single Cell Controller (10-fold genomics) system in the NCI-CCR single-cell analysis tool. Dead cells were removed, the cells were adjusted to the required concentration, and approximately 8000 cells were detected on the machine. Sequencing was performed on Nova Seq (Illumina) in the NCI-CCR sequencing facility. Bioinformatics analysis was performed using the OmicStudio tools at https://www.omicstudio.cn/tool.

### Statistical Analyses

All data in the present study are presented as the mean ± standard deviation. The significance of the differences between two groups was calculated using a two-tailed unpaired Student’s t-test. Analysis of variance and Tukey’s post-hoc tests were performed for more than two groups (multiple comparisons). All statistical analyses were performed using GraphPrism (v8.0). Statistical significance was set at P < 0.05. All intensities of fluorescence expression in the experiments were calculated using the ImageJ software. The standard symbols were presented as *P < 0.05, **P < 0.01, ***P < 0.005, and ****P < 0.001.

## Acknowledgment

This work was supported by National Natural Science Foundation of China (No. 32022043). This work is partly supported by Collaborative Innovation Center of Suzhou Nano Science & Technology, the Priority Academic Program Development of Jiangsu Higher Education Institutions (PAPD), the 111 Project.

## Competing interests

The authors declare no competing interests.

## Author contributions

C.W. and F. L. designed the project. Q. Y. and H. D. performed the experiments and collected the data. Q. Y. and H. D. analyzed and interpreted the data. All authors contributed to the writing of the manuscript, discussed the results and implications, and edited the manuscript at all stages.

## Data Transparency Statement

data, analytic methods, and study materials will be made available to other researchers.

---

**Fig. S1.**
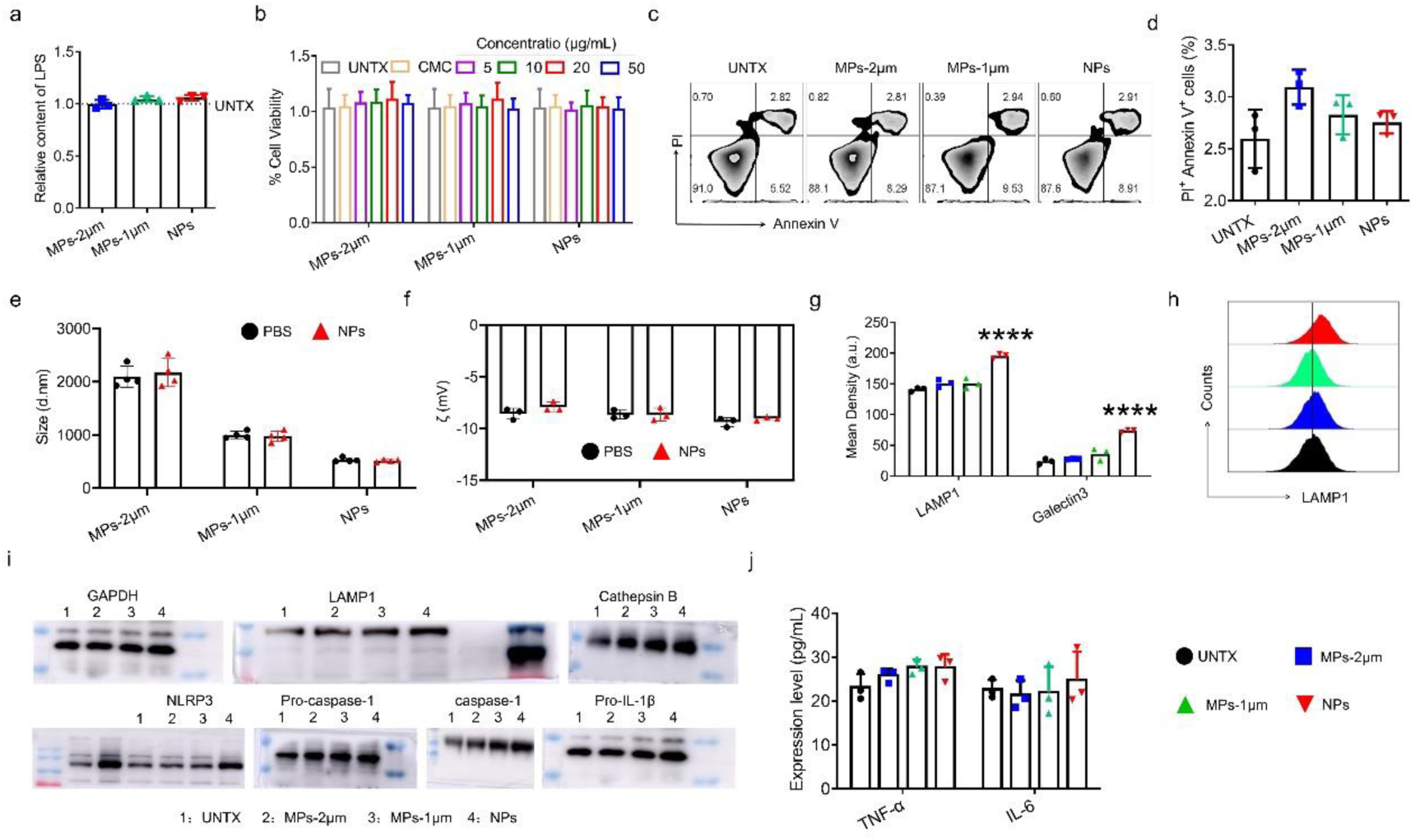
Cytotoxicity of MPs and NPs and lysosomal damage to macrophages. (**a**) Detection of LPS in cell supernatant after treatment. (**b**) The MTT assay curves of Raw264.7 cells for 24 h of NPs and MPs treatment. (**c-d**) Apoptosis detection by Annexin V/ PI double staining of Raw264.7 cells for 24 h of NPs and MPs treatment. (**e**) Particle size distribution measured by DLS and (**f**) zeta potential in PBS (pH=7) after co-incubated with cell culture medium. (**g**) Quantification of mean fluorescence intensity (MFI) of LAMP1 and galectin3. (**h**) Flow cytometric analysis of LAMP1 in Raw264.7 cells (gated on F4/80^+^). (**i**) Western blot analysis of the expression of various types of proteins in RAW264.7 cells. (**j**) Cytokine analysis of IL-6 and TNF-α in cell supernatant. Data are shown as mean ± SD (n=3-4). Statistical significance was calculated by one-way ANOVA using the Tukey posttest. \****P*** < 0.05; \*\****P*** < 0.01; \*\*\****P*** < 0.005; \*\*\*\****P*** < 0.001. a. u., arbitrary units.

**Fig. S2.**
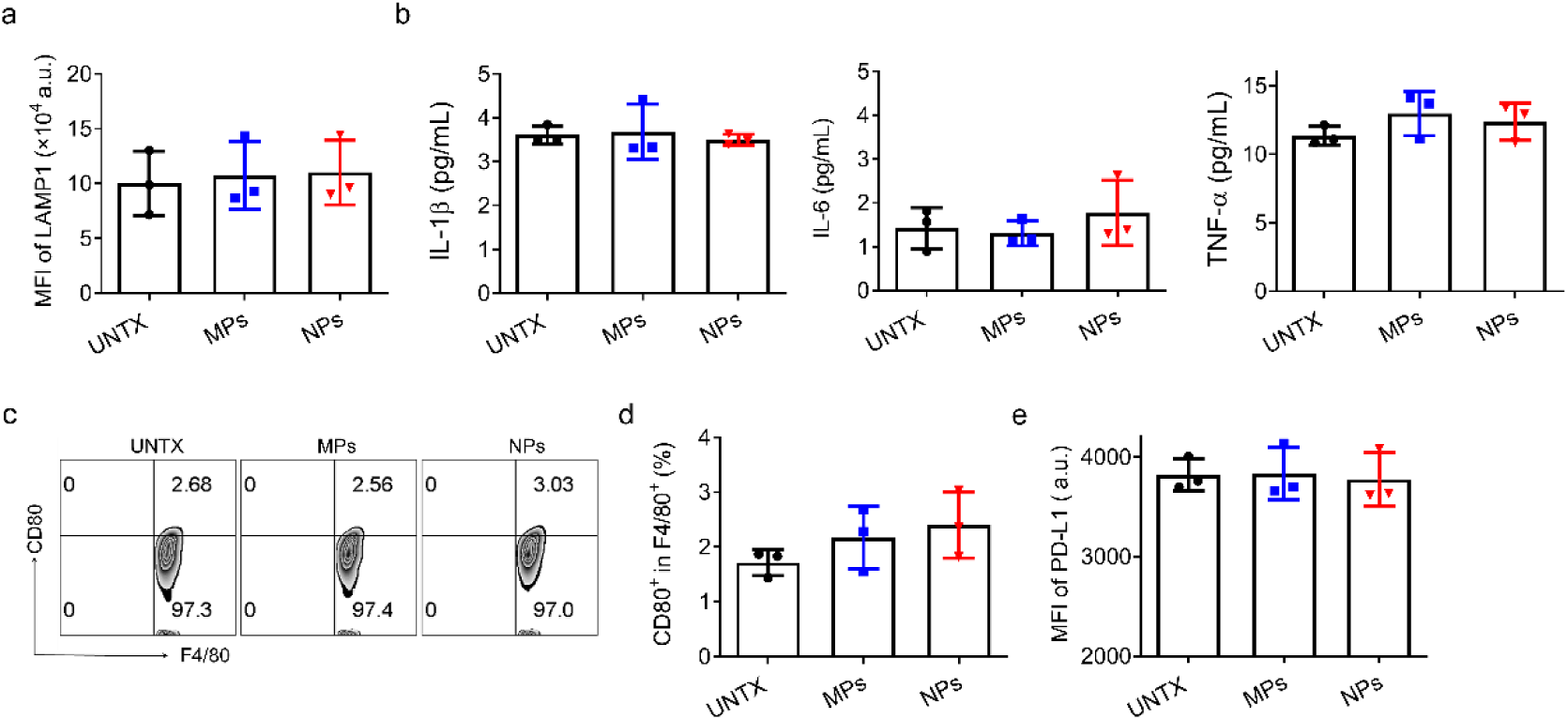
Cytochalasin B inhibits the effect of NPs. (**a**) Flow cytometric quantification of the MFI of LAMP1. (**b**) Cytokine analysis of IL-1β, IL-6 and TNF-α in cell supernatant. (**c**) Flow cytometric analysis of CD80 expression on Raw264.7 cells and (**d**) corresponding quantitative analysis. (**e**) The expression of PD-L1 on Raw264.7 cells. Data are shown as mean ± SEM (n=3). Statistical significance was calculated by one-way ANOVA using the Tukey posttest. \****P*** < 0.05; \*\****P*** < 0.01; \*\*\****P*** < 0.005; \*\*\*\****P*** < 0.001. a. u., arbitrary units.

**Fig. S3.**
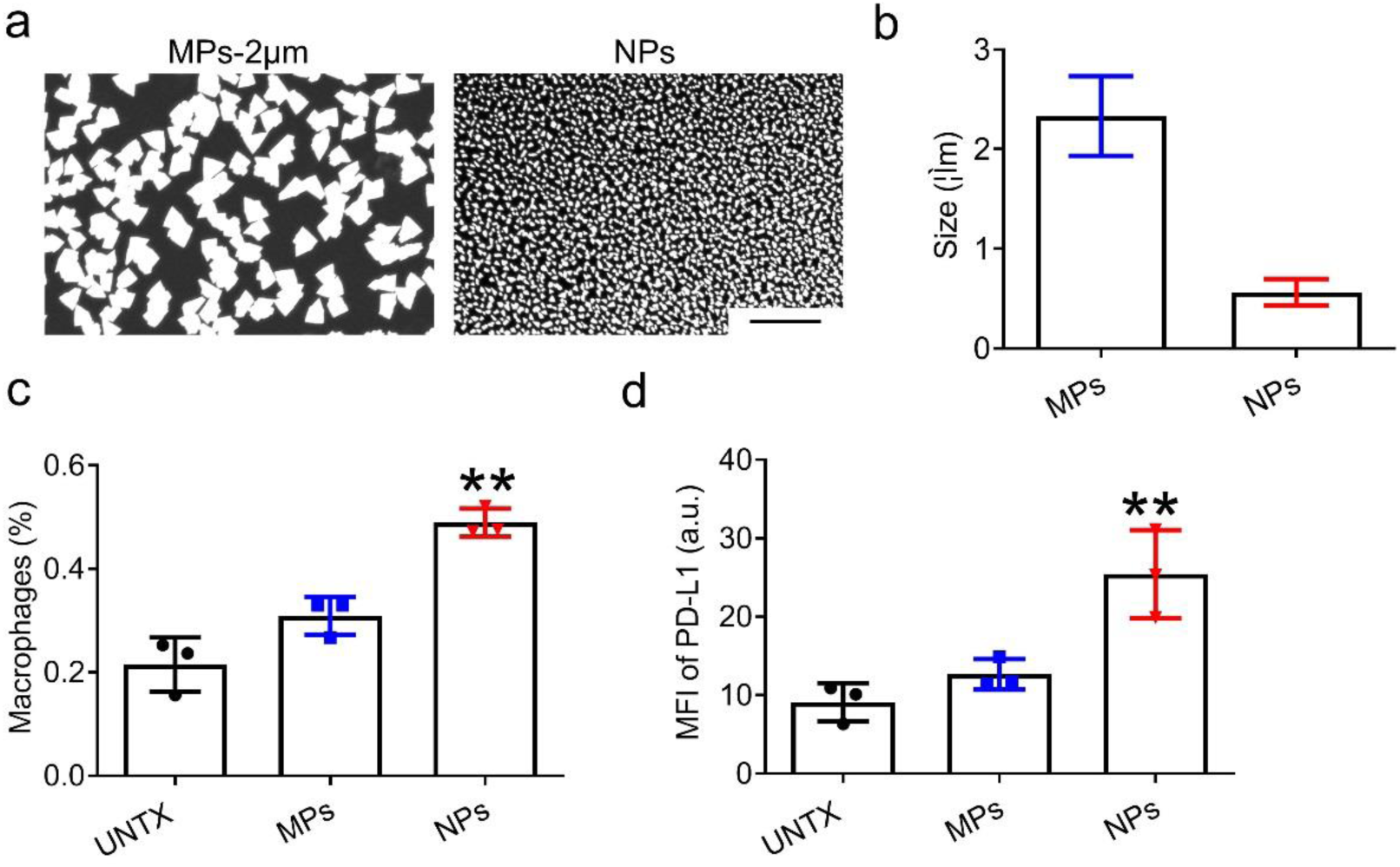
MPs and NPs are not degraded by gastric fluid and NPs induce intestinal pro-inflammatory macrophages. (**a**) SEM images of MPs and NPs after treatment with simulated gastric fluid and (**b**) size statistics. Scale bar: 10μm. (**c**) Ratio of intestinal macrophages (F4/80^+^). (**d**) Expression of PD-L1 on intestinal macrophages after MPs and NPs treatment. Data are shown as mean ± SD (n=3). Statistical significance was calculated by one-way ANOVA using the Tukey posttest. \****P*** < 0.05; \*\****P*** < 0.01; \*\*\****P*** < 0.005; \*\*\*\****P*** < 0.001. a. u., arbitrary units.

**Fig. S4.**
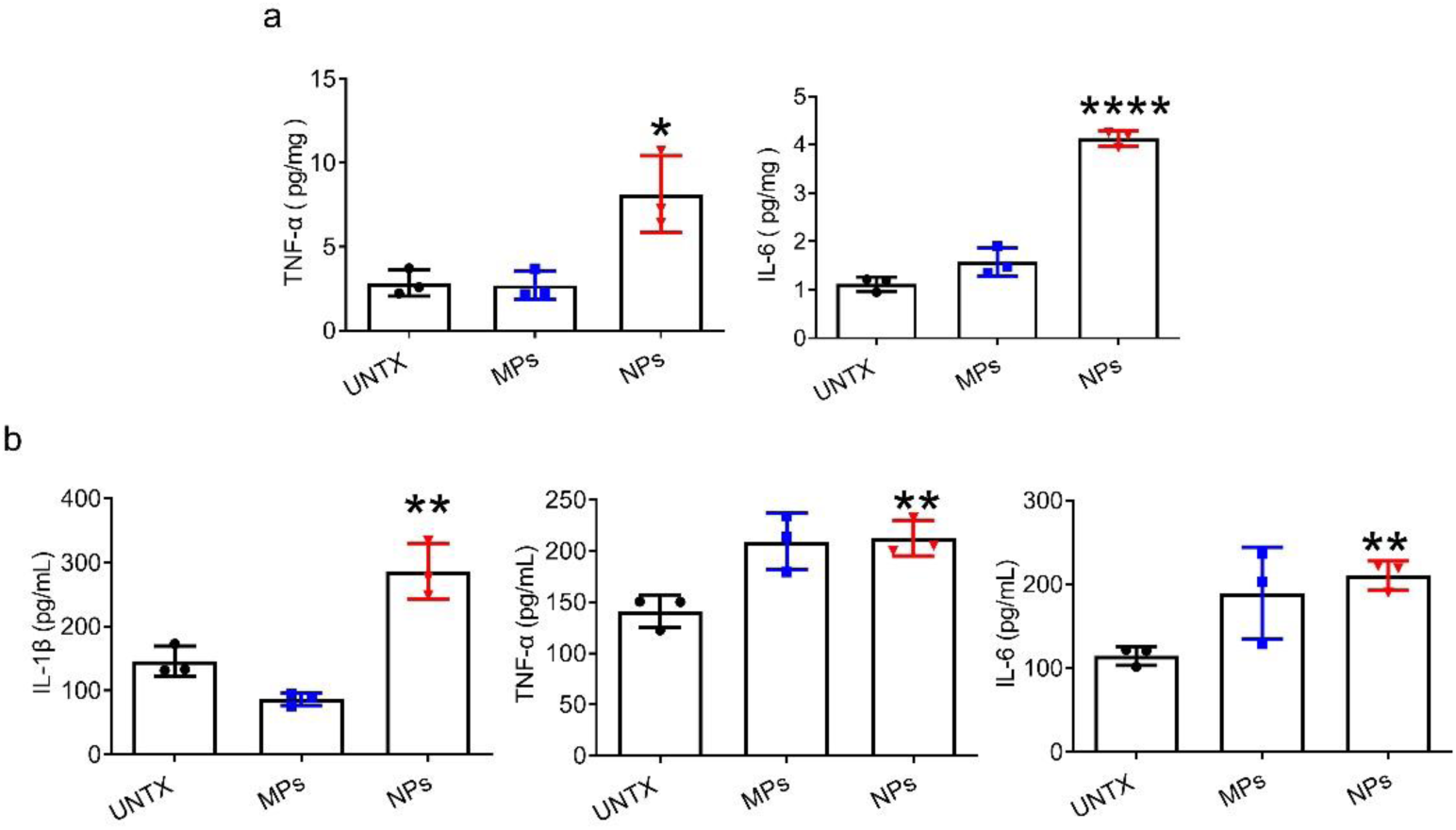
NPs increase the levels of pro-inflammatory cytokines in the intestine and circulation. (**a**) Analysis of cytokine content in intestine and (**b**) in serum after MPs and NPs treatment. Data are shown as mean ± SD (n=3). Statistical significance was calculated by one-way ANOVA using the Tukey posttest. \****P*** < 0.05; \*\****P*** < 0.01; \*\*\****P*** < 0.005; \*\*\*\****P*** < 0.001.

**Fig. S5.**
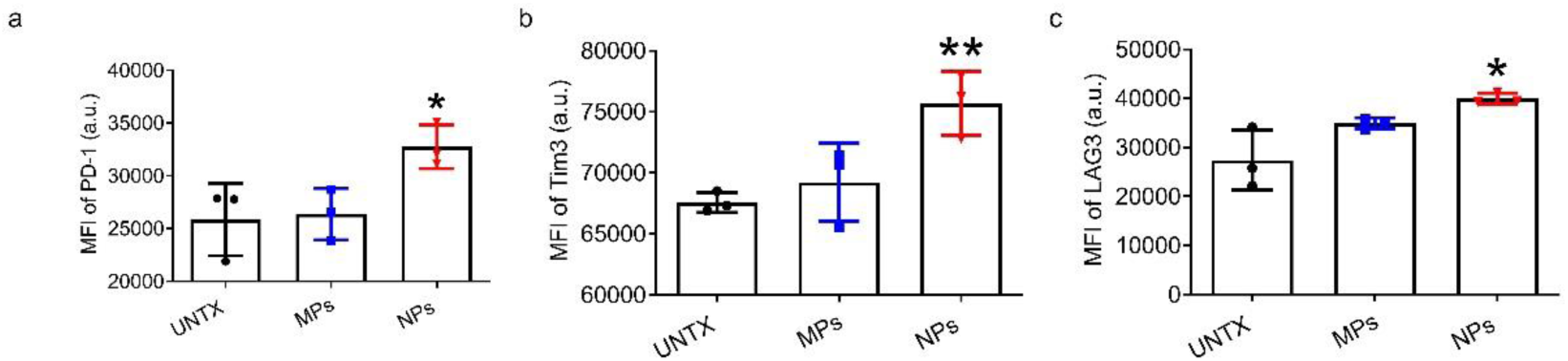
NPs lead to dysfunctional phenotype of T cell. (**a**) Flow cytometry analysis of PD-1 in CD3^+^ T cells. (**b**) Flow cytometry analysis of TIM3 and (**c**) LAG3 in CD3^+^ T cells. Data are shown as mean ± SD (n=3). Statistical significance was calculated by one-way ANOVA using the Tukey posttest. \****P*** < 0.05; \*\****P*** < 0.01; \*\*\****P*** < 0.005; \*\*\*\****P*** < 0.001. a. u., arbitrary units.

**Fig. S6.**
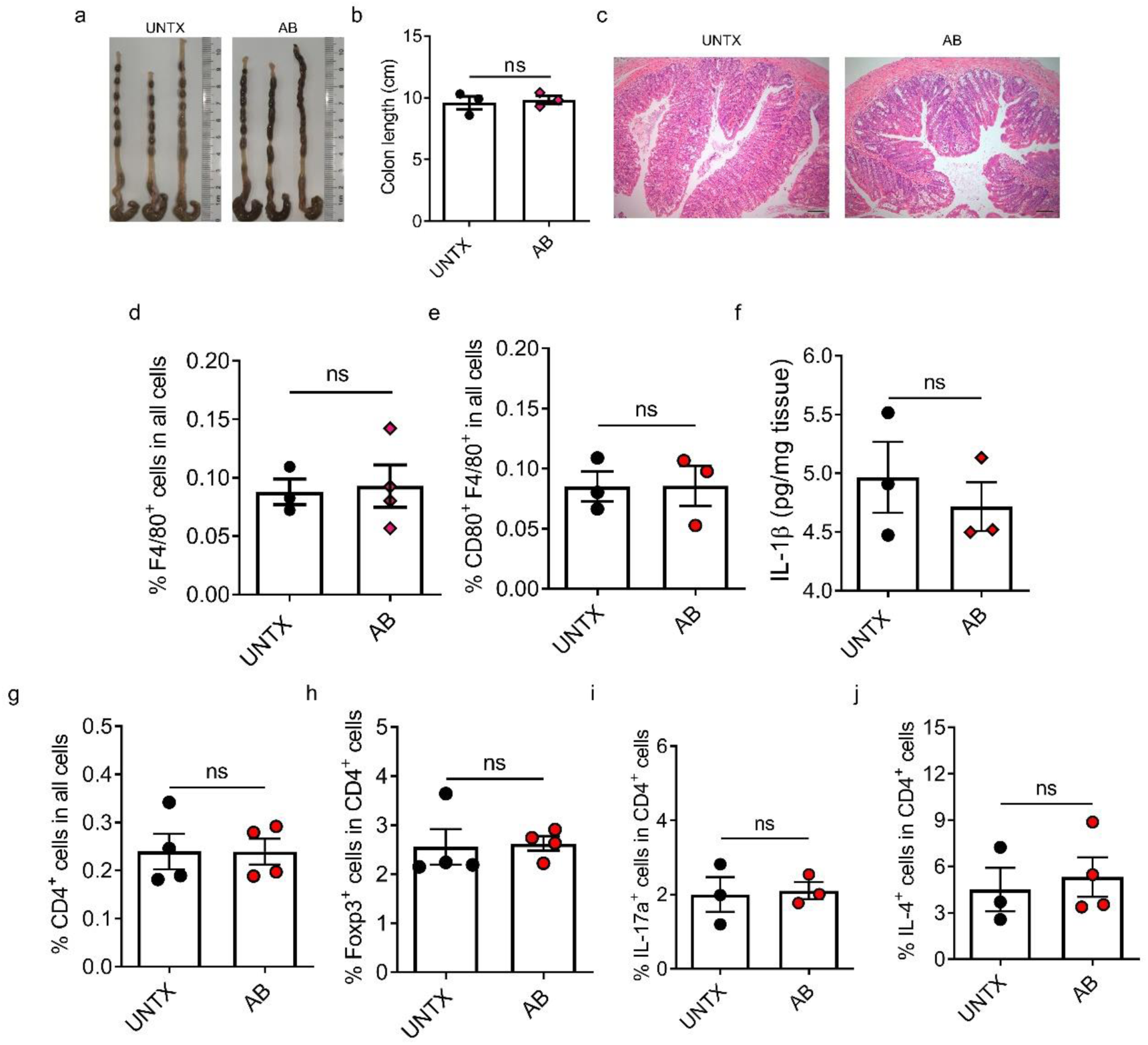
Antibiotic cocktails therapy showed no obvious effect on intestinal macrophage and CD4^+^ T cells in our experiment. (**a**) Representative images of the colon of mice fed with three different sizes of plastic particles. (**b**) Quantification of the colon length. (**c**) Representative images of H&E-stained colon sections illustrate inflammatory cell infiltrates. Scale bars, 100 μm. (**d**) The percentages of F4/80^+^ cells in all cells. (**e**) Flow cytometric analysis of CD80 expression on F4/80^+^ cells. (**f**) The level of IL-1β in the intestine measured by ELISA. (**g**) The percentages of CD4^+^ cells in all cells and Foxp3^+^ (**h**), IL-17a^+^ (**i**), and IL-4^+^ (**j**) in CD4^+^ T cells in the mice fed with different particle. Data are shown as mean ± SD (n=3-4). Statistical significance was calculated by Student’s t-test (two-tailed). *P < 0.05; **P < 0.01; ***P < 0.005; ****P < 0.001.

**Fig. S7.**
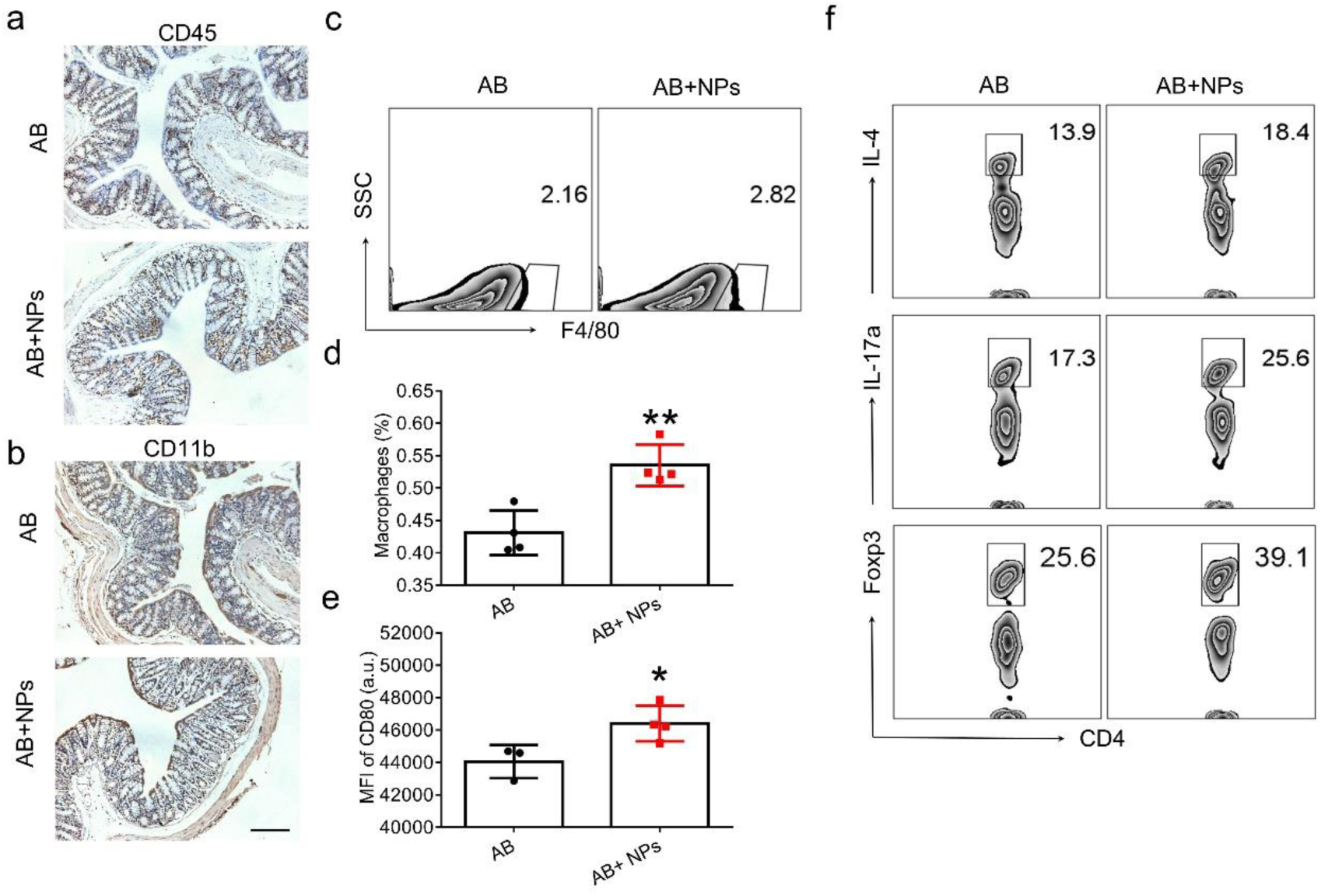
NPs cause an increase in proinflammatory macrophages and differentiation of CD4^+^ T cells after antibiotic treatment. (**a-b**) Immunohistochemical imaging of CD45 and CD11b in mice treated with antibiotic before NPs treatment. (**c**) Flow cytometry analysis of macrophages and (**d**) corresponding quantification. (**e**) Detection of CD80 on macrophages. (**f**) Flow cytometry analysis of CD4^+^ T cells in intestine. Data are shown as mean ± SD (n=3-4). Statistical significance was calculated by Student’s t-test (two-tailed). \****P*** < 0.05; \*\****P*** < 0.01; \*\*\****P*** < 0.005; \*\*\*\****P*** < 0.001.

**Fig. S8.**
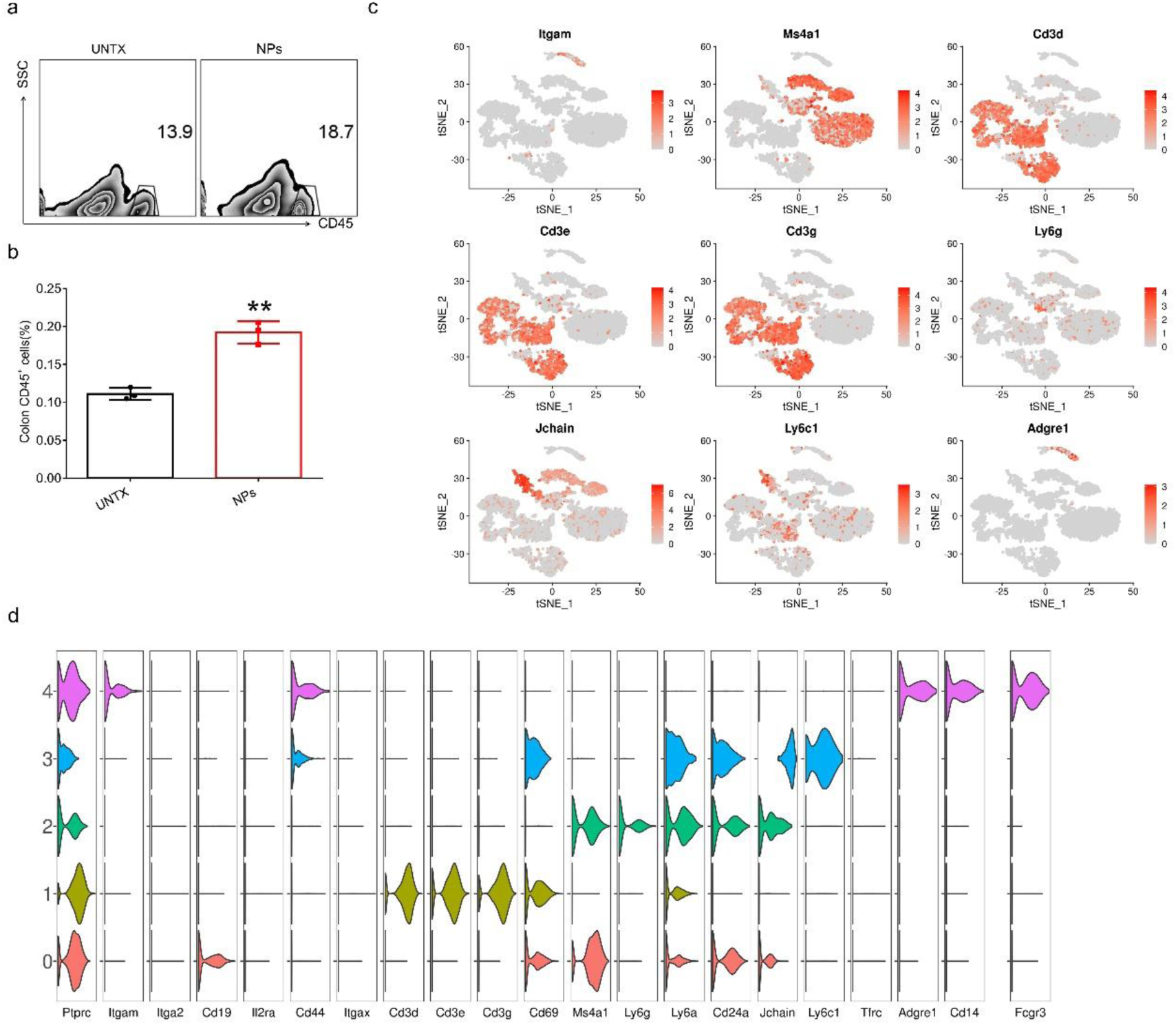
NPs increase the proportion of intestinal CD45^+^ cells. (**a**) Flow cytometric analysis of intestinal CD45^+^ cells and (**b**) corresponding quantitative analysis. (**c**) tSNE map of marker genes expression. (**d**) Violin diagram of gene expression. Data are shown as mean ± SD (n=3). Statistical significance was calculated by Student’s t-test (two-tailed). \****P*** < 0.05; \*\****P*** < 0.01; \*\*\****P*** < 0.005; \*\*\*\****P*** < 0.001.

**Fig. S9.**
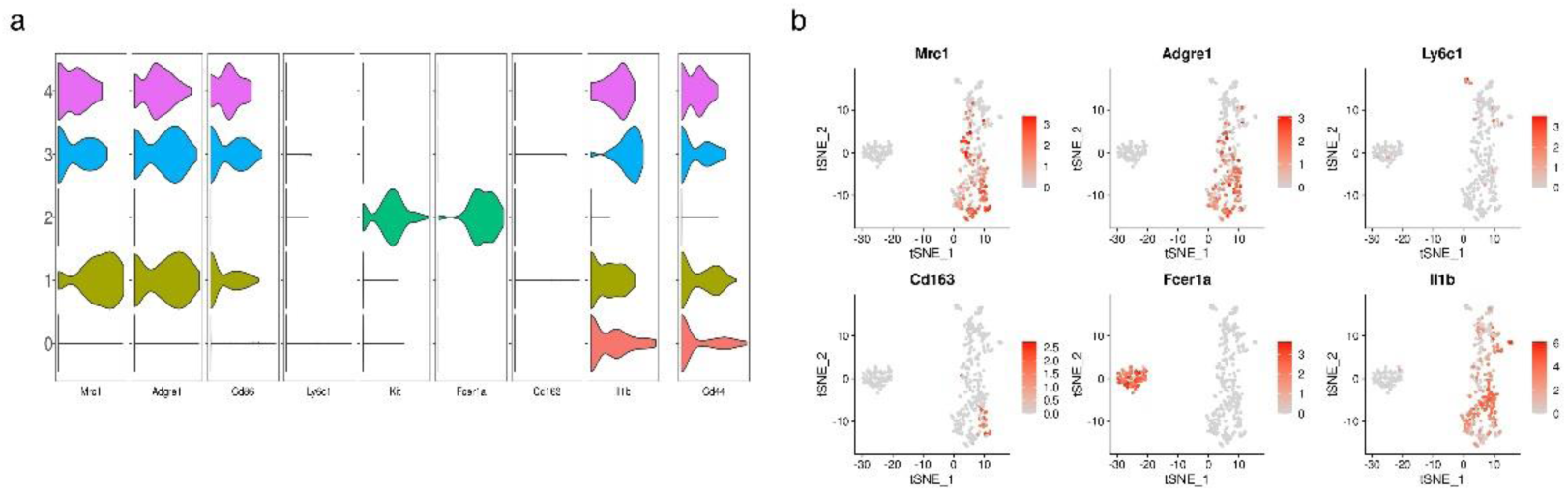
NPs reduce intestinal mast cells. (**a**) Violin diagram of gene expression in MΦ. (**b**) tSNE map of marker genes expression. C: UNTX; T: NPs.

**Fig. S10.**
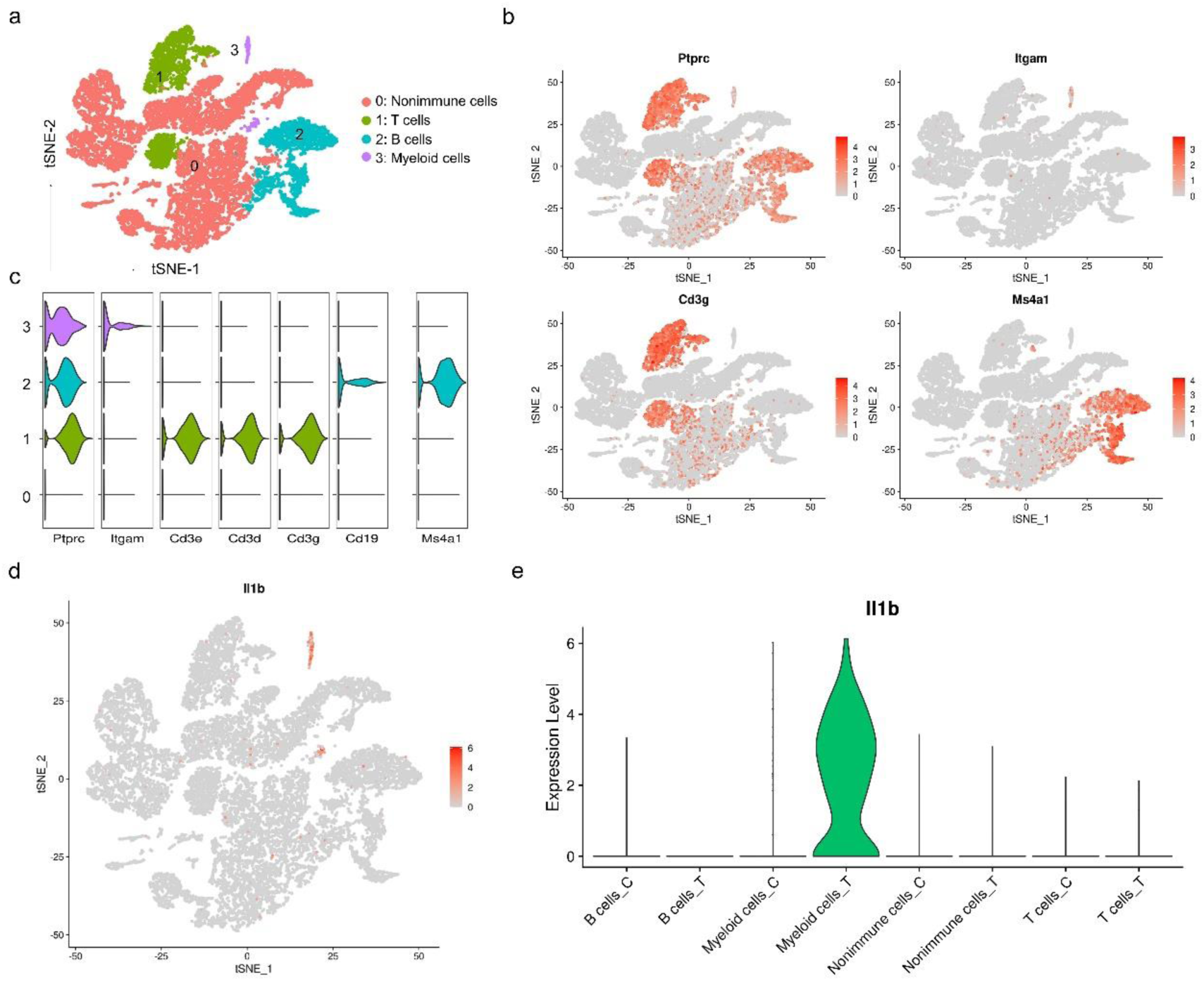
NPs upregulates *Il1b* gene expression in intestinal myeloid cells. (a) tSNE image of all intestinal cells. (b) tSNE map and (c) violin diagram of marker genes expression in intestinal cells. (d) tSNE map and (c) violin diagram of *Il1b* expression in intestinal cells. C: UNTX, T: NPs.

**Fig. S11.**
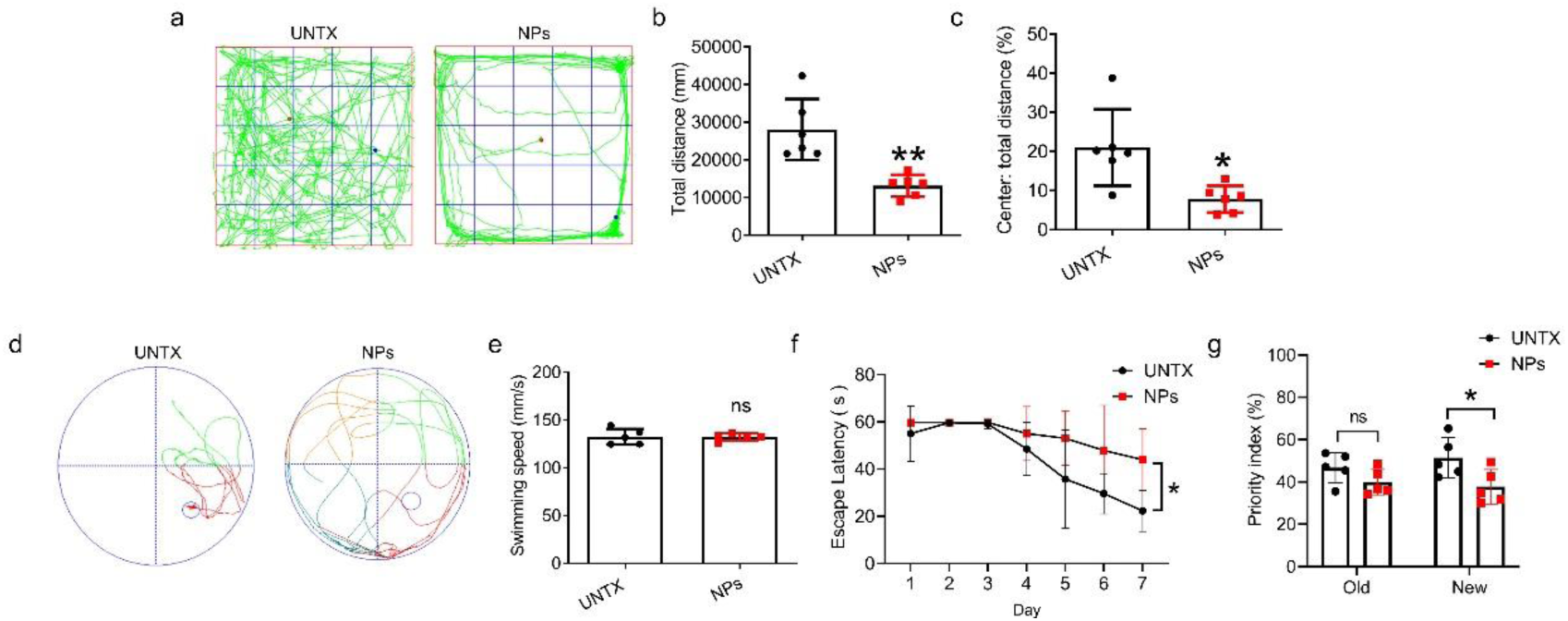
NPs exacerbate cognitive and memory dysfunction in APP/PS1 (AD) mice after 2-months treatment. (**a**) Open field test images and (**b**) quantitative analysis of the total distance and (**c**) the ratio of the center distance to the total distance. (**d**) Plots of Morris water maze images and quantification of (**e**) the average swimming speed and (**f**)the escape latency during day 1 to day 7. (**g**) Recognition index in the Novel Object Recognition. Data are shown as mean ± SD (n=4-6). Statistical significance was calculated by Student’s t-test (two-tailed) and one-way ANOVA using the Tukey posttest. \****P*** < 0.05; \*\****P*** < 0.01; \*\*\****P*** < 0.005; \*\*\*\****P*** < 0.001. a. u., arbitrary units.

**Fig. S12.**
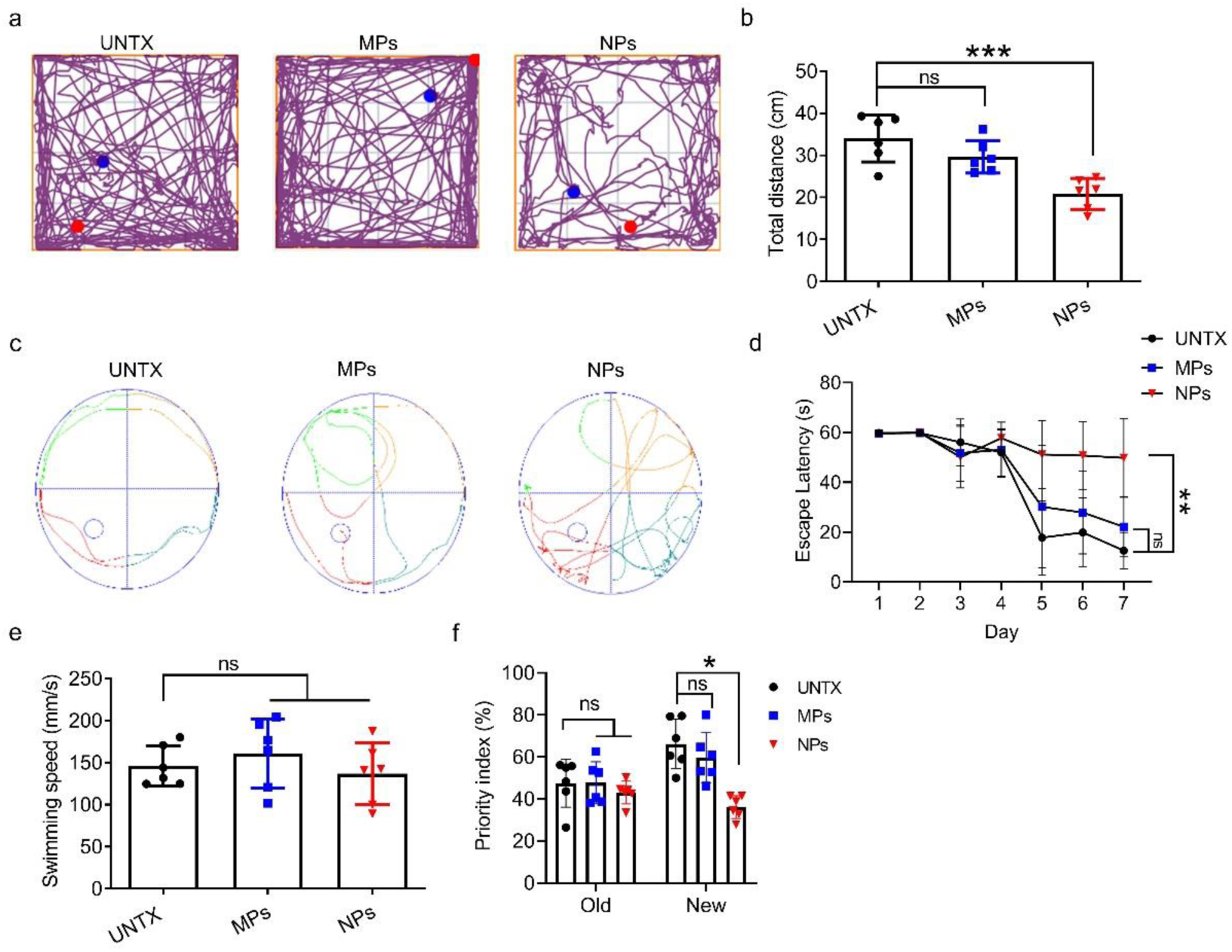
NPs are more likely than MPs to cause cognitive and memory dysfunction in mice. (**a**) Representative open field test images and (**b**) quantitative analysis of the total distance. (**c**) Plots of Morris water maze images and quantification of (d) escape latency of day1 to day 7 and (**e**) the average swimming speed. (**f**) Recognition index in the novel object recognition (NOR). Data are shown as mean ± SD (n=6). Statistical significance was calculated by one-way ANOVA using the Tukey posttest. \****P*** < 0.05; \*\****P*** < 0.01; \*\*\****P*** < 0.005; \*\*\*\****P*** < 0.001.

**Fig. S13.**
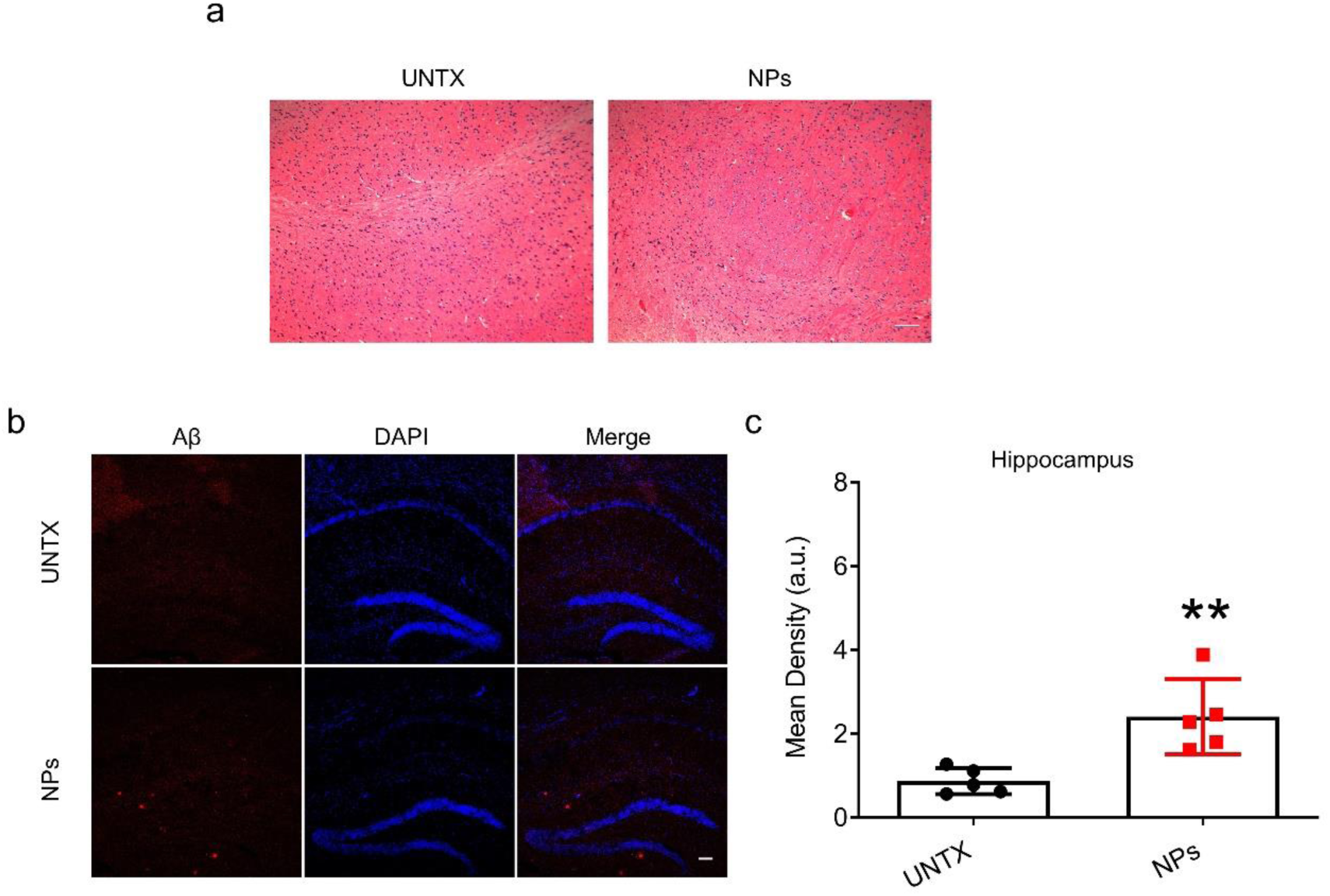
NPs induce the elevation of the level of Aβ in hippocampus. (**a**) H&E images of the brain. Scale bars, 100μm. (**b**) Confocal microscopy images of Aβ protein in hippocampus and (**c**) mean fluorescence quantification. Data are shown as mean ± SD (n=5). Statistical significance was calculated by Student’s t-test (two-tailed) using the Tukey posttest. \****P*** < 0.05; \*\****P*** < 0.01; \*\*\****P*** < 0.005; \*\*\*\****P*** < 0.001. a. u., arbitrary units.

**Fig. S14.**
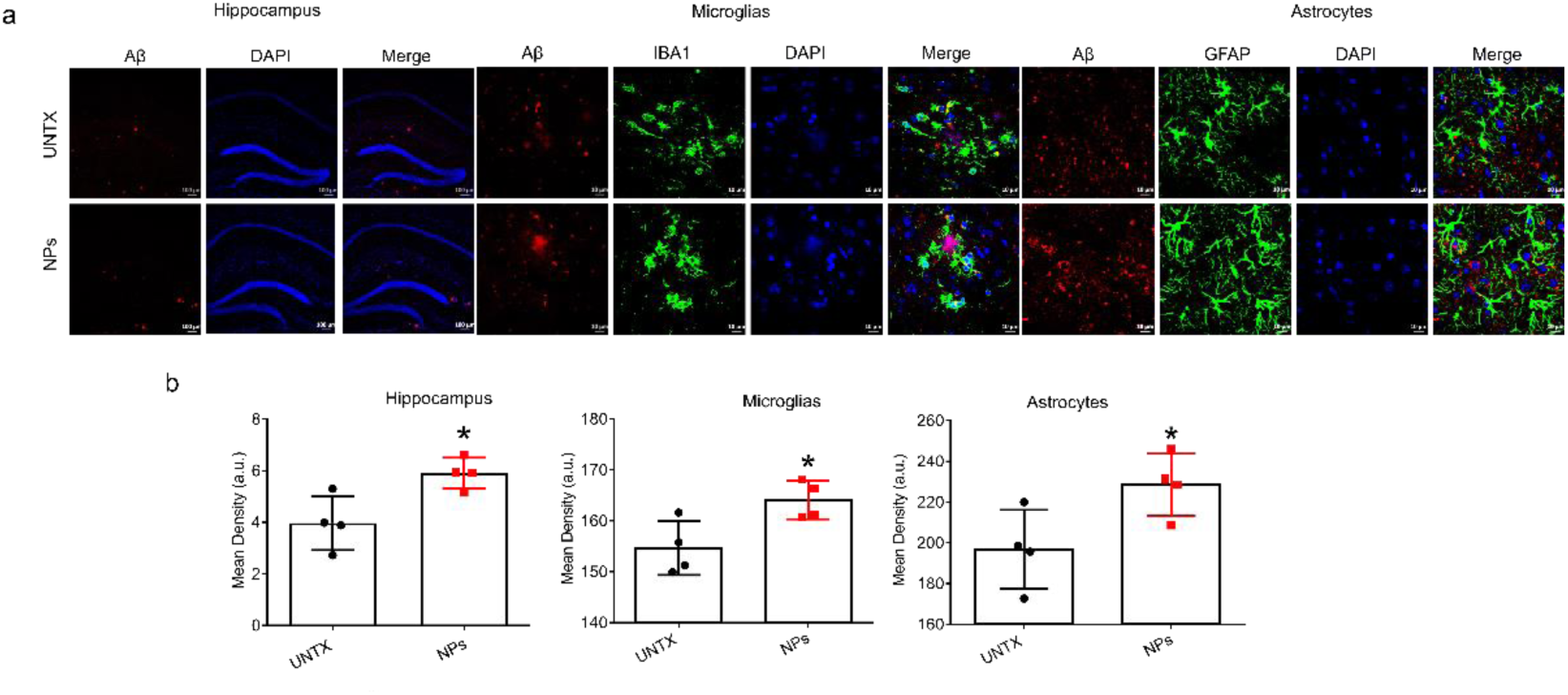
(a) Confocal microscopy images of Aβ protein in hippocampus, astrocytes and microglia and (b) mean fluorescence quantification of Aβ protein. Data are shown as mean ± SD (n=4-6). Statistical significance was calculated by Student’s t-test (two-tailed) and one-way ANOVA using the Tukey posttest. *P < 0.05; **P < 0.01; ***P < 0.005; ****P < 0.001. a. u., arbitrary units.

**Fig. S15.**
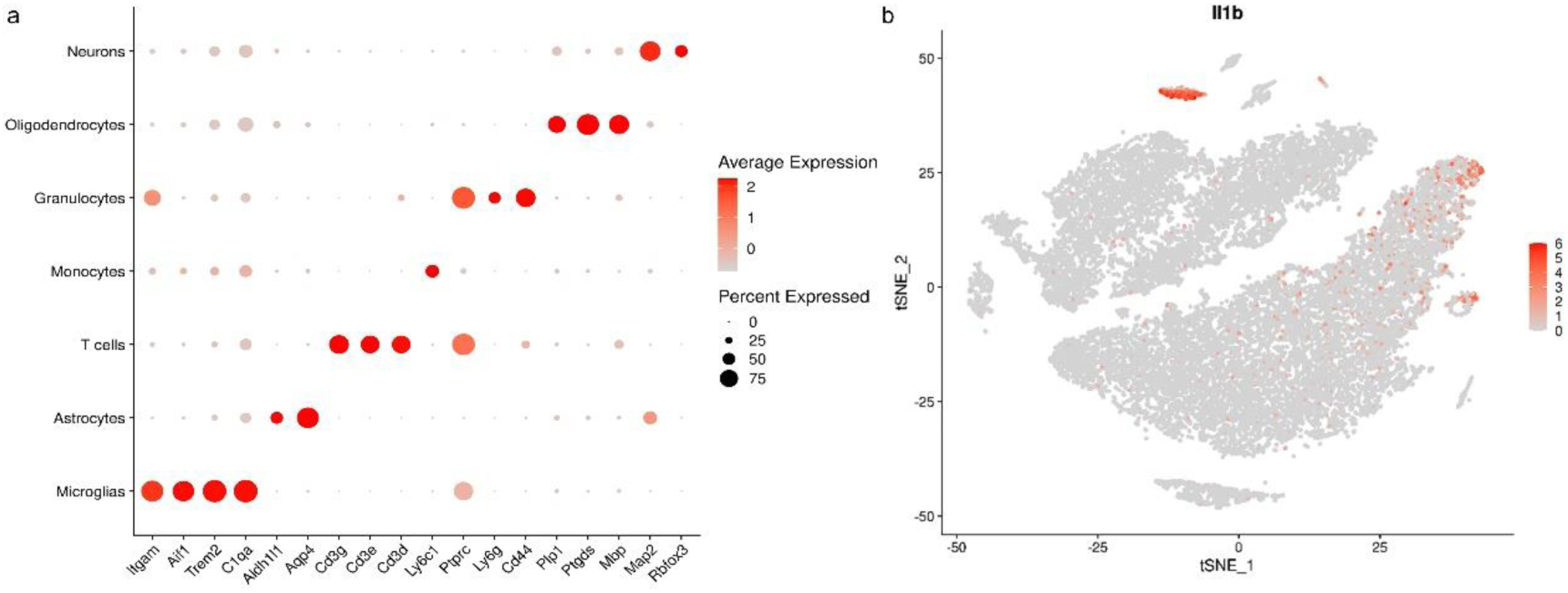
NPs do not alter *Il1b* expression in the brain cells. (a) Dotplot of marker genes expression in all brain cell clusters. (b) t-SNE map of *Il1b* expression in brain cells.

**Fig. S16.**
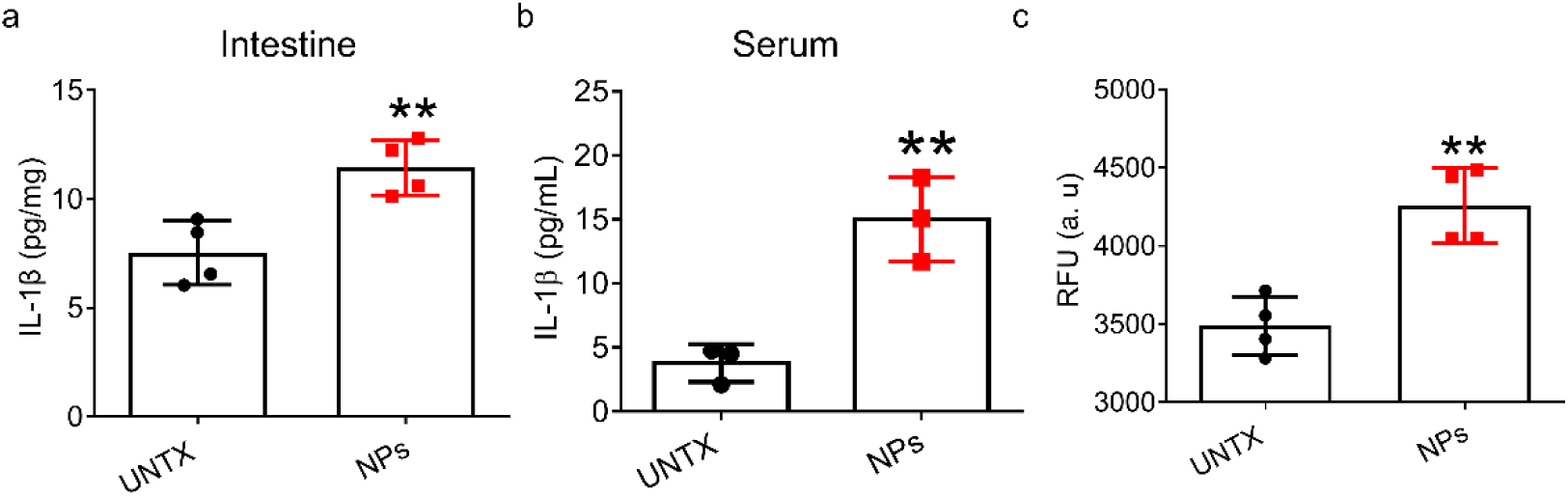
Long-term uptake of NPs increases IL-1β levels in the intestine, circulation, and brain. (**a**) Analysis of cytokine detection in the intestine. (**b**) Analysis of cytokine detection in the serum. (**c**) The relative fluorescence intensity of FITC-dextran in serum. RFU: relative fluorescence units. Data are shown as mean ± SD (n=4). Statistical significance was calculated by Student’s t-test (two-tailed). \****P*** < 0.05; \*\****P*** < 0.01; \*\*\****P*** < 0.005; \*\*\*\****P*** < 0.001. a. u., arbitrary units.

**Fig. S17.**
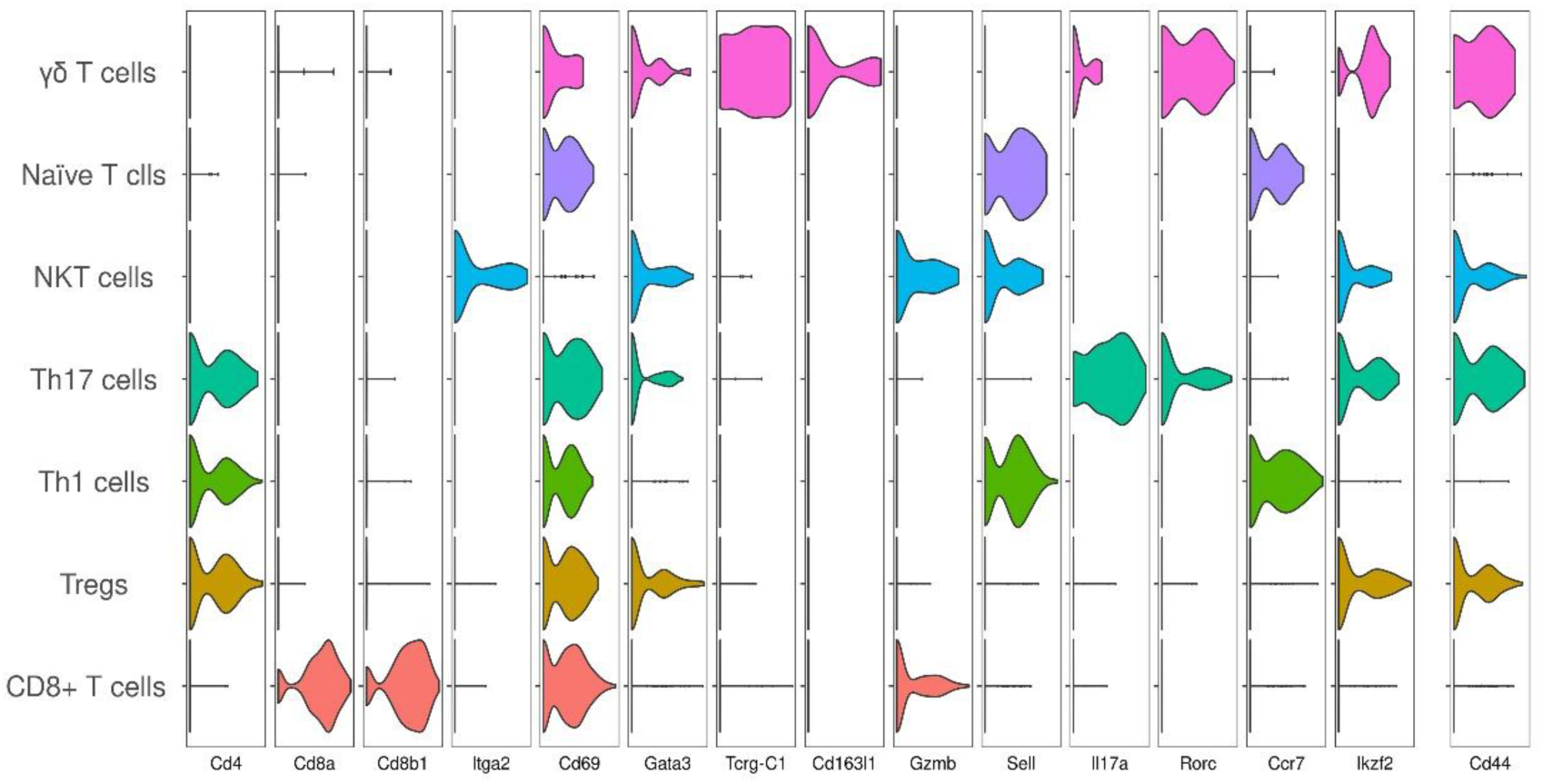
Violin diagram of marker genes expression in clusters of T cells.

**Fig. S18.**
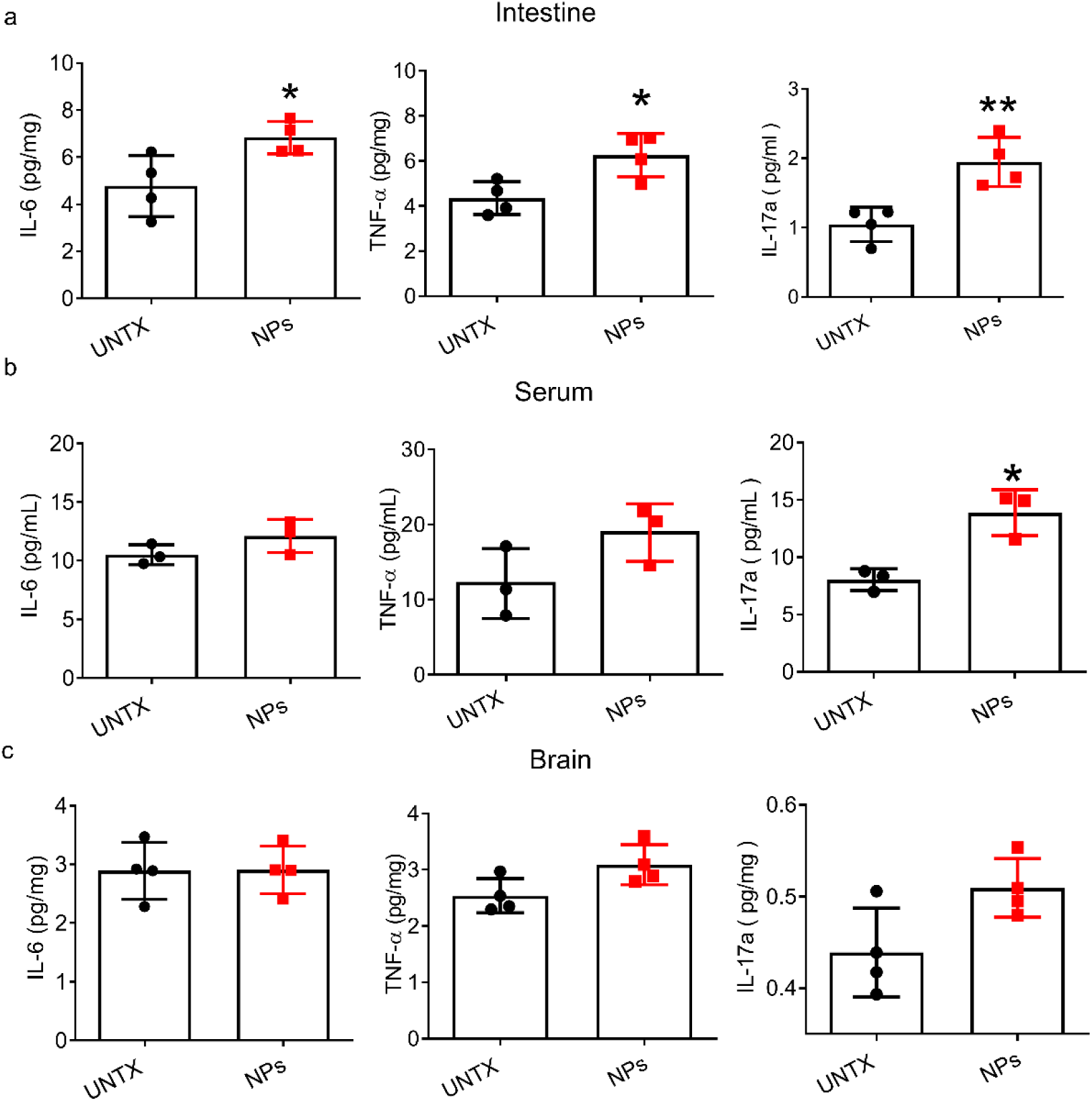
Long-term uptake of NPs increases IL-17a levels in the intestine, circulation, and brain. (**a**) Analysis of cytokine detection in the intestine. (**b**) Analysis of cytokine detection in the serum. (**c**) Analysis of cytokine detection in the brain. Data are shown as mean ± SD (n=4). Statistical significance was calculated by Student’s t-test (two-tailed). \****P*** < 0.05; \*\****P*** < 0.01; \*\*\****P*** < 0.005; \*\*\*\****P*** < 0.001. a. u., arbitrary units.

**Fig. S19.**
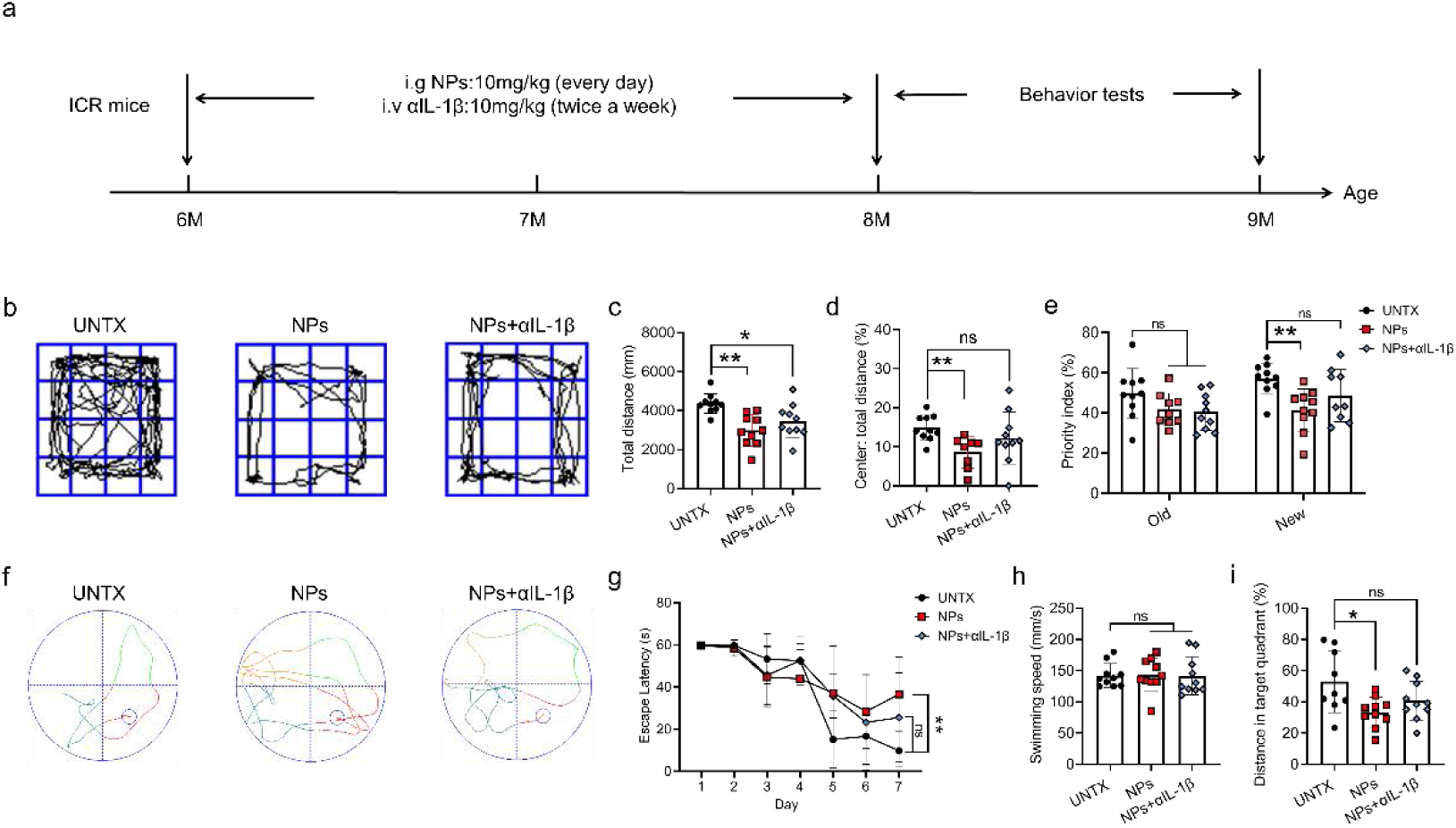
Neutralizing IL-1β produced by NPs treatment alleviates cognitive and memory impairment in ICR mice. (**a**) Experimental timeline. (**b**) Representative open field test images and (**c**) quantitative analysis of the total distance and (**d**) the ratio of the center distance to the total distance. (**e**) Recognition index in the novel object recognition (NOR). (**f**) Plots of Morris water maze images and quantification of (**g**) escape latency, (**h**) swimming speed, and (**i**) the ratio of the distance in the target quadrant to the total distance. Data are shown as mean ± SD (n=10). Statistical significance was calculated by one-way ANOVA using the Tukey posttest. \****P*** < 0.05; \*\****P*** < 0.01; \*\*\****P*** < 0.005; \*\*\*\****P*** < 0.001.

**Fig. S20.**
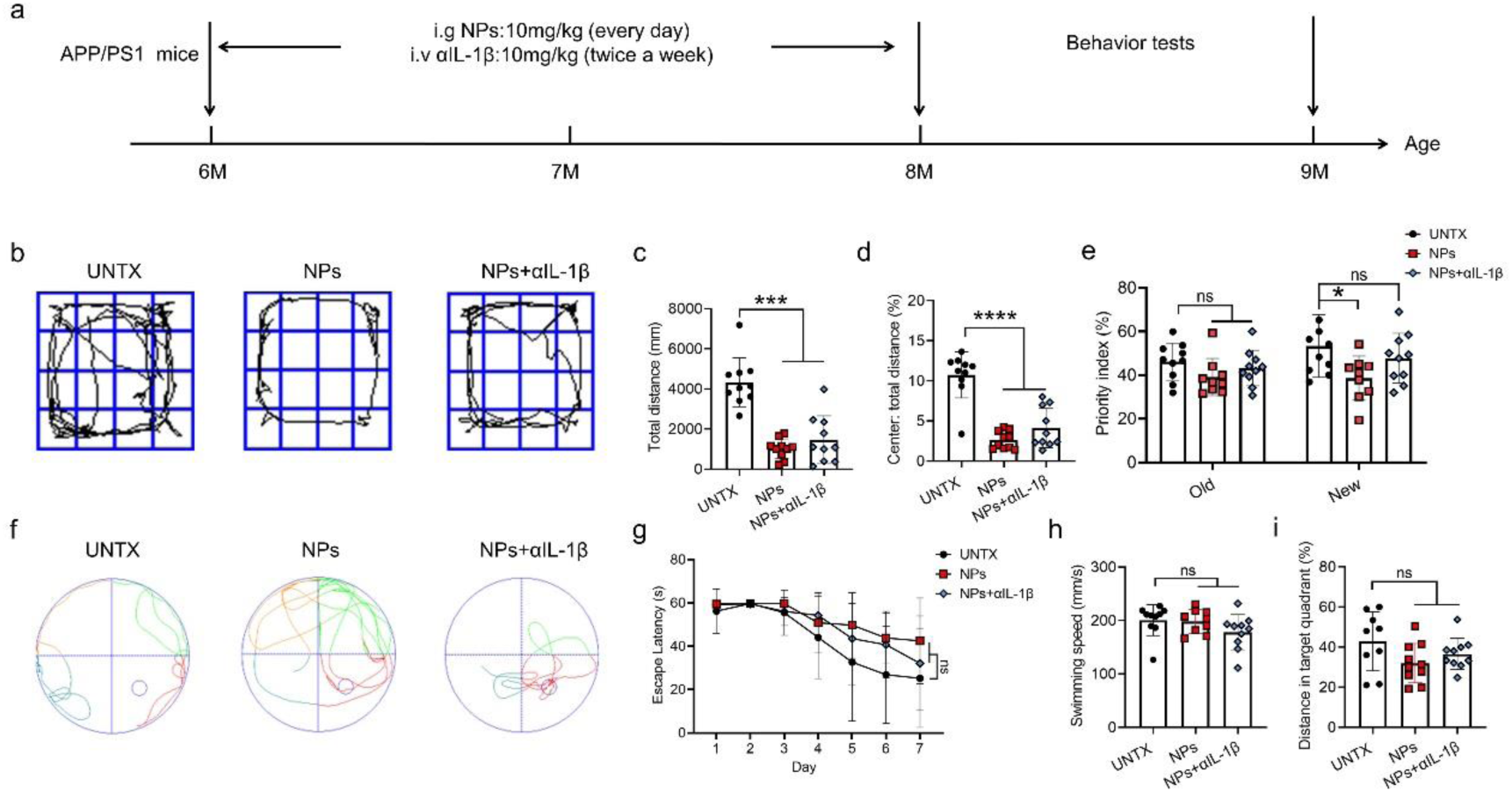
Neutralizing IL-1β in APP/PS1 NP-fed mice. (**a**) Experimental timeline. (**b**) Representative open field test images and (**c**) quantitative analysis of the total distance and (**d**) the ratio of the center distance to the total distance. (**e**) Recognition index in the novel object recognition (NOR). (**f**)Plots of Morris water maze images and quantification of (**g**) escape latency, (**h**) swimming speed, and (**i**) the ratio of the distance in the target quadrant to the total distance. Data are shown as mean ± SD (n=10). Statistical significance was calculated by one-way ANOVA using the Tukey posttest. \****P*** < 0.05; \*\****P*** < 0.01; \*\*\****P*** < 0.005; \*\*\*\****P*** < 0.001.

**Fig. S21.**
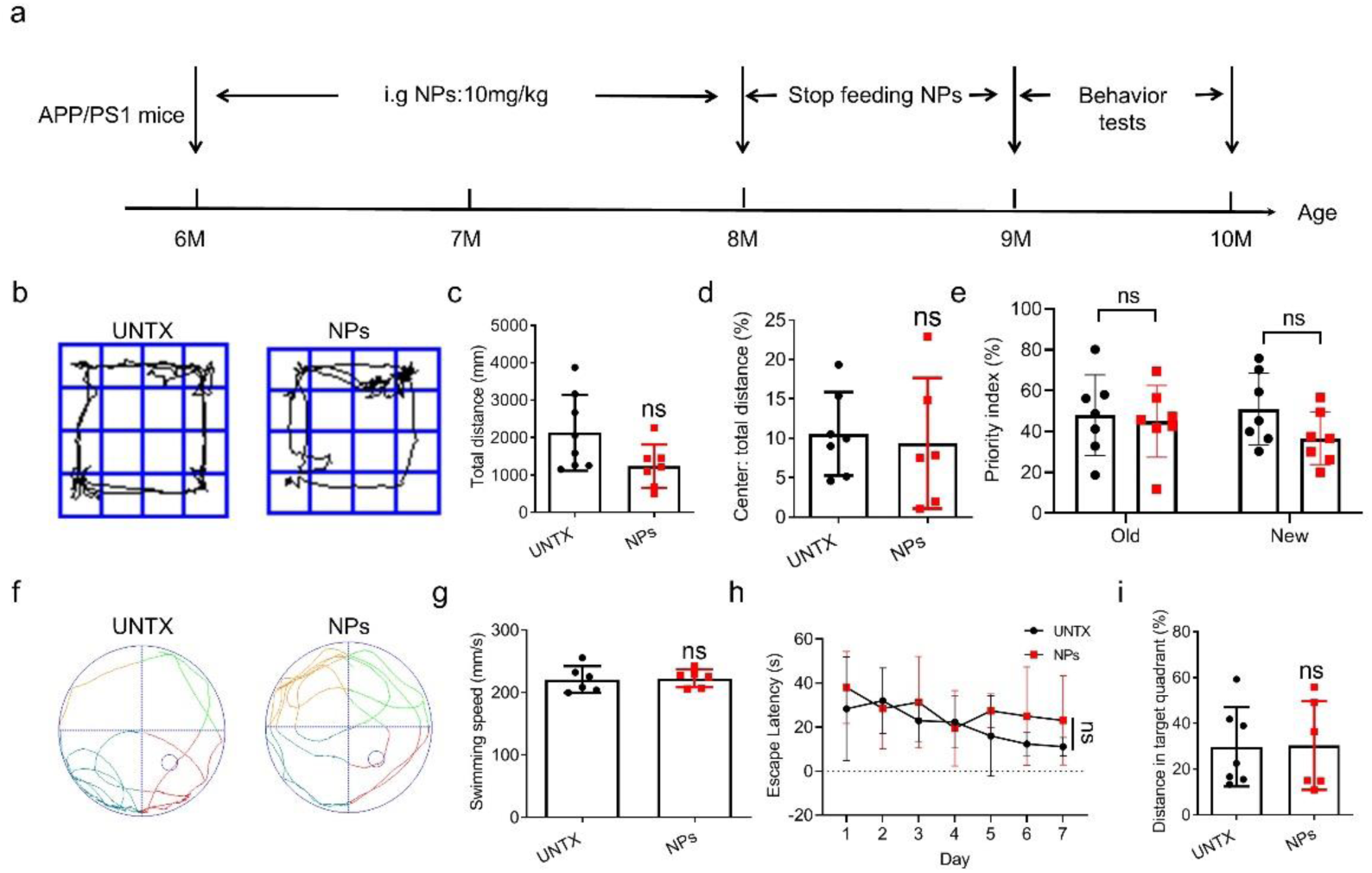
Discontinuation of NPs feeding alleviates cognitive and memory dysfunction in APP/PS1 mice treated with NPs for two months. (a) Experimental timeline. (b) Representative open field test images and (c) quantitative analysis of the total distance and (d) the ratio of the center distance to the total distance. (e) Recognition index in the novel object recognition (NOR). (f)Plots of Morris water maze images and quantification of (g) escape latency, (h) swimming speed, and (i) the ratio of the distance in the target quadrant to the total distance. Data are shown as mean ± SD (n=6-8). Statistical significance was calculated by one-way ANOVA using the Tukey posttest. \**P* < 0.05; \*\**P* < 0.01; \*\*\**P* < 0.005; \*\*\*\**P* < 0.001.

## Notes

### Competing Interest Statement

The authors have declared no competing interest.

